# Diverse task-driven modeling of macaque V4 reveals functional specialization towards semantic tasks

**DOI:** 10.1101/2022.05.18.492503

**Authors:** Santiago A. Cadena, Konstantin F. Willeke, Kelli Restivo, George Denfield, Fabian H. Sinz, Matthias Bethge, Andreas S. Tolias, Alexander S. Ecker

**Author notes:** Equal senior author contributions (Fabian H. Sinz, Matthias Bethge, Andreas S. Tolias, Alexander S. Ecker).

## Abstract

Responses to natural stimuli in area V4 – a mid-level area of the visual ventral stream – are well predicted by features from convolutional neural networks (CNNs) trained on image classification. This result has been taken as evidence for the functional role of V4 in object classification. However, we currently do not know if and to what extent V4 plays a role in solving *other* computational objectives. Here, we investigated normative accounts of V4 (and V1 for comparison) by predicting macaque single-neuron responses to natural images from the representations extracted by 23 CNNs trained on different computer vision tasks including semantic, geometric, 2D, and 3D types of tasks. We found that V4 was best predicted by semantic classification features and exhibited high task selectivity, while the choice of task was less consequential to V1 performance. Consistent with traditional characterizations of V4 function that show its high-dimensional tuning to various 2D and 3D stimulus directions, we found that diverse non-semantic tasks explained aspects of V4 function beyond those captured by individual semantic tasks. Nevertheless, jointly considering the features of a pair of semantic classification tasks was sufficient to yield one of our top V4 models, solidifying V4’s main functional role in semantic processing and suggesting that V4’s affinity to 2D or 3D stimulus properties found by electrophysiologists can result from semantic functional goals.

## Introduction

What is the functional role of area V4 in primate visual information processing? One line of evidence suggests that V4 is tuned in a high-dimensional space that facilitates the joint encoding of shape and surface characteristics of object parts [1, 2] (e.g. sensitivity to luminance [3], texture [4] and chromatic contrasts [5], blurry boundaries [6], and luminance gradients [7]). Although very insightful, these experiments are constrained to a relatively small number of stimulus feature directions that potentially miss other important aspects of V4 function that could be unlocked with richer natural stimuli. Recent work used a *transfer learning* approach to infer the functional role of different brain areas. The features extracted by convolutional neural networks (CNNs) pre-trained on object classification *transfer* well to the task of predicting V4 responses to natural stimuli [8–11]. This result has been interpreted as evidence that object recognition is one of the major goals of V4 processing. However, a natural question arises: Do other computational goals beyond object classification explain V4 responses equally well or even better? Recent work using fMRI in humans has attempted to assign different functional goals to different regions of interest in the brain [12–14], but it remains unclear whether single neurons express the same patterns of selectivity as the highly aggregated, indirect fMRI signal.

Inferring the functional role of a brain area using transfer learning is a promising avenue, but it is complicated by the fact that transfer performance is not only determined by the pre-training task itself, but also by the size of the dataset used for pre-training, the network architecture and other factors. A recent development in the computer vision community could be very promising for the neuroscience community, because it mitigates some of these problems: The *taskonomy* project [15] released a dataset that consists of 4.5 million images, ground truth labels for 23 different visual tasks for each image, and pre-trained convolutional neural networks with the same architecture (ResNet50) on each task.

We employed the *taskonomy* project to investigate how well the representations learned by training on each of these visual tasks predict single-cell responses to natural images recorded in macaque areas V4 and V1 (Fig. 1). Using this approach, we can isolate the contribution of different pre-training tasks on how well the learned representations match those of areas V1 and V4 without the results being confounded by different network architectures or datasets across tasks. We found that a diverse set of tasks explained V1 responses almost equally well, while scene and object classification tasks provided better accounts for V4 responses than all the alternative tasks tested. We further built models that jointly read from pairs of task representations and found that 2D, 3D, and geometric types of tasks capture additional nonlinearities beyond those captured by individual semantic tasks, consistent with descriptions of V4’s high-dimensional tuning observed by electrophysiologists [1, 16].

**Figure 1.**
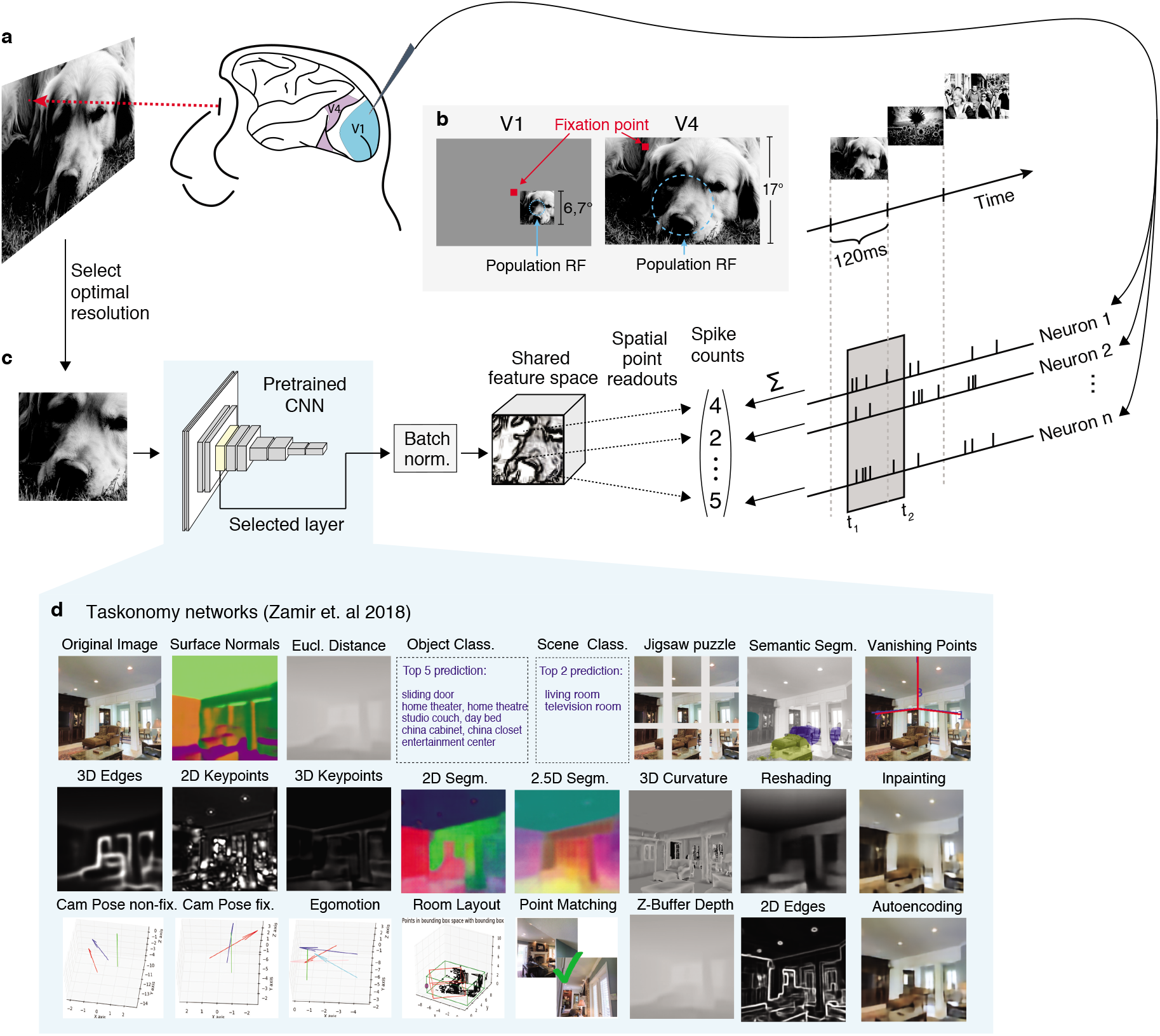
Experimental paradigm and diverse task modeling of neural responses. **A**, natural images were shown in sequence to two fixating rhesus macaques for 120ms while neural activity was recorded with a laminar silicon probe. Following careful spike-sorting, spike counts were extracted in time windows 40–160ms (V1) and 70–160ms (V4) after image onset. The screen covered *∼* 17*^◦^ ×* 30*^◦^* of visual field with a resolution of *∼* 63px*/^◦^*. For each area, we showed approx. 10k unique images once (*train set*). A random set of 75 images, identical for both areas, was repeated 40-55 times (*test set*). **B**, For V1 recordings (left), the fixation spot was centered on the screen. Square, gray-scale, 420px ImageNet images (6.7*^◦^*) were placed at the center of each session’s population receptive field (size: *∼* 2*^◦^*, eccentricities: 2*^◦^ −* 3*^◦^*). In the V4 recordings sessions (right), the fixation spot was accommodated to bring the population receptive field as close to the center of the screen as possible (size: *∼* 8*^◦^*, eccentricities: 8*^◦^ −* 12*^◦^*). All images were up-sampled and cropped to cover the whole screen. We isolated 458 (V1) and 255 (V4) neurons from 32 sessions of each area. **C**, Predictive model. Cropped input images covering 2.7*^◦^*(V1) and 12*^◦^* (V4) were resized and forwarded through the first *l* layers of a pretrained convolutional neural network (CNN) to produce features that are then batch-normalized and shared by all neurons. The input scale factor was a hyper-parameter, cross-validated on a held-out subset of the train set (*validation set*). The point readout [17] extracts features at a single spatial location and computes a regularized linear mapping to the neural responses for each neuron separately (see Methods). The readout and batch-normalization parameters were jointly learned to minimize the Poisson loss between predicted and observed response rates. **D**, *taskonomy* networks used for feature extraction (Adapted with permission from Zamir et al. (2018) [15]). We used the pretrained encoder CNNs of these tasks, which share a Resnet50 architecture [18], to build our models and compare their predictive abilities on V1 and V4.

However, combining the features of both object and scene classification was sufficient to obtain peak V4 performance, indicating that multiple semantic goals can induce complementary intermediate representations that are predictive of the additional nonlinearities contributed by non-semantic tasks. Overall, our results solidify V4’s semantic functional role and explain that V4’s affinity to other non-semantic tasks can result from semantic computational goals.

## Results

We collected datasets of well-isolated single-cell responses from V4 and primary visual cortex (V1) for comparison. We measured the spiking activity of individual neurons from two awake, fixating rhesus macaques (M1, M2) using a 32-channel linear array spanning multiple cortical layers [19, 20], in response to tens of thousands of grayscale natural images presented in sequence over many trials (Fig. 1a). These images were sampled uniformly from the ImageNet Large-Scale Visual Recognition Challenge (ILSVRC-2012) dataset [21] and displayed for 120 ms each without interleaving blanks (see Methods). Most of these images were shown only once (*train-set*) while a selection of 75 images was repeated multiple times (*test-set*). We isolated 458 V1 neurons from 15 (M1) and 17 (M2) sessions at eccentricities 2–3°; and 255 V4 neurons from 11 (M1) and 21 (M2) sessions at eccentricities 8–12°. For the V1 sessions, we centered the stimuli on the population receptive field of the neurons. For the V4 sessions, the stimuli covered the entire screen. We obtained image–response pairs by extracting spike counts in the windows 40–160 ms (V1) and 70–160 ms (V4) after image onset (Fig. 1a–c), which corresponded to the typical response latency of the neurons in the respective brain area. We computed the stimulus-driven variability of spike counts using the repeated trial presentations on the test-set (see Methods) and found that the mean [*±* s.d.] fraction of explainable variance of these two areas was not significantly different (0.31 *±* 0.18 (V1) and 0.32 *±* 0.19 (V4); two-sided *t* -test, *t*(711) = *−*1.152*, p* = 0.2). Following previous work [20], we excluded unreliable neurons from the performance evaluations where the fraction of explainable variance was lower than 0.15, yielding 202 (V4) and 342 (V1) neurons (Fig. S2a).

### Task-driven modelling of neural responses

We built upon the *taskonomy* project [15], a recent large-scale effort of the computer vision community, in which CNN architectures consisting of *encoder-decoder* parts were trained to solve various visual tasks. The *encoder* provides a low-dimensional representation of the input images from which each task can be (nonlinearly) read-out by the *decoder*. We considered the encoder network of 23 of these tasks, which have been previously categorized into *semantic*, *geometric*, *2D*, and *3D* groups (listed in Fig. 1d) based on hierarchical clustering of their encoder representations [15]. We chose these networks because of two key features: 1) all of them were trained on the same set of images, and 2) all encoder networks have the same architecture (ResNet-50 [18]). Any differences we observe across the learned representations are thus caused by the training objective targeted to solve a specific task.

To quantify the match between the representations extracted by intermediate layers of the *taskonomy* networks and V4 representations, we used these networks to built task-driven models [20, 22] of single-neuron recordings in response to natural stimuli: We presented the images that were shown to the monkey to each pretrained network, then extracted the resulting output feature maps from several intermediate layers and fed these to a regularized linear-nonlinear (LN) readout that was specific to each recorded neuron (Fig. 1c). This readout acted on the features at a single spatial location [17], preventing any additional nonlinear spatial integration beyond what has been computed by the task-trained network. For each *taskonomy* network, we built one model for each readout layer and optimized regularization parameters for each model. To ensure that the resulting correspondence between network layers and neural data is not merely driven by the confound between growing receptive field sizes and feature complexity along the network’s depth, we optimized the resolution of the input images on held-out data from the training set (Supplementary Fig. S1). This prevented us from assigning V4 to a layer simply because of the matching receptive field coverage that could result from an arbitrary input resolution, but instead allowed us to find for each model the layer with the best aligned nonlinearities to the data.

### V1 is better predicted by task representations than V4

We evaluated the predictive performance of our fitted task-driven models with the correlation between model predictions and the average response across trials. When accounting for input scale (Supplementary S1), we found that performance on V4 peaks at a higher layer than V1 (Fig. 2a–b, Supplementary Fig. S3, a signature of hierarchical correspondence and increased complexity of V4 over V1 found in anatomical and latency studies [23]. This functional hierarchy was present in most cases, but could not be explained by the architecture alone: V1 and V4 were assigned to the same layer for networks trained on texture edges, jigsaw, vanishing points, and the untrained baseline (a network with matching architecture and random weights). However, for several 2D tasks, the hierarchical assignment was not so strict as higher layers in the network retained high predictive power in comparison to other tasks (Fig. S3).

**Figure 2.**
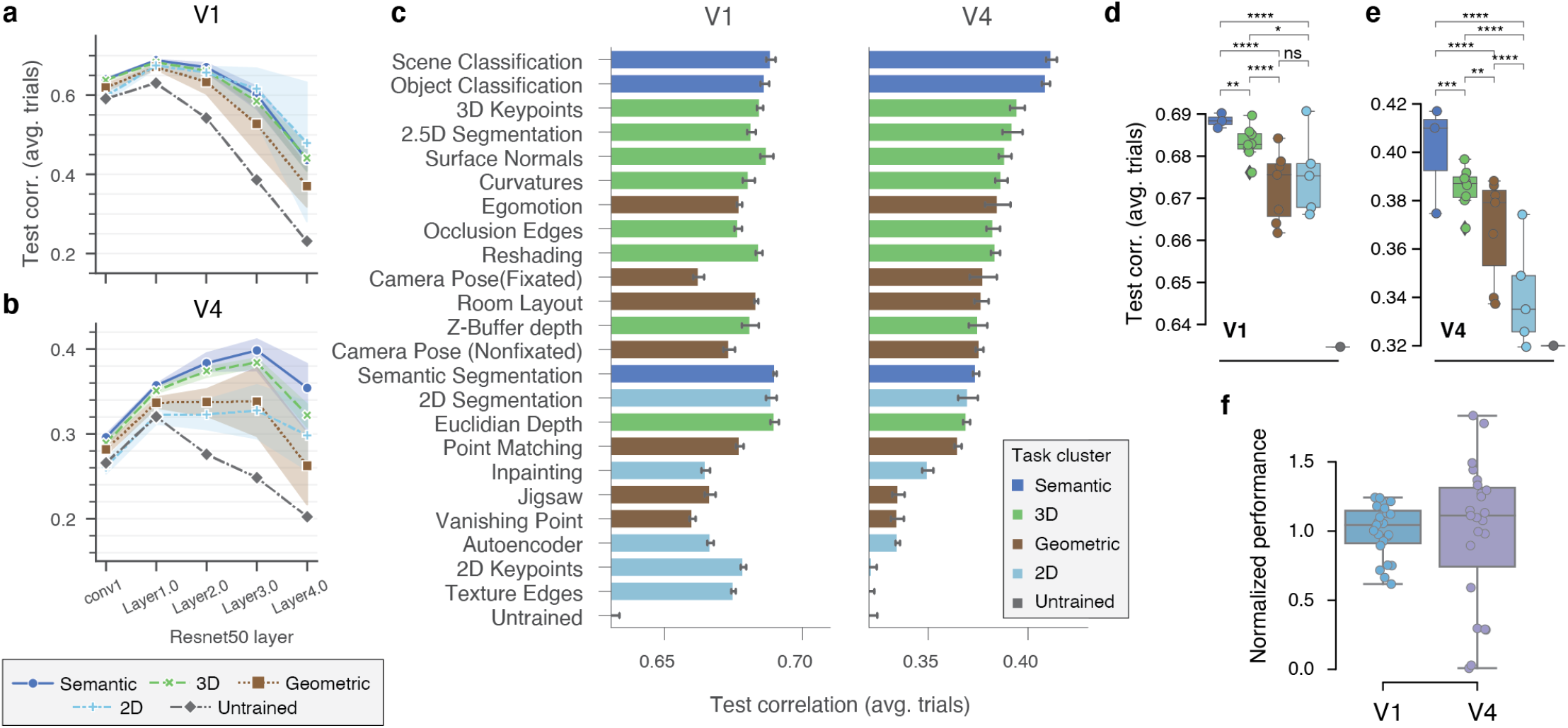
Comparison of diverse task-driven models on V1 and V4. **a,b**, Average performance of task clusters (see Fig. 1f) on V1 (**a**) and V4 (**b**) as a function of network layer. Error bands represent 95% confidence intervals across individual tasks (see Supplementary Fig. S3). Performance is measured as the average test-set correlation across neurons between model predictions and mean spike counts over repeated presentations. All tasks outperform an untrained, random baseline that shares the same architecture (dark gray). The best predictive features for V1 can be found at an early layer (Layer1.0) for all models, while deeper layers yielded peak performance for V4 with semantic and 3D task-clusters outperforming the others. **c**, The individual task-model performance on V1 (left) and V4 (right), maximized over layers and other hyper-parameters in the validation set. Error bars show 1 s.e. of the mean for five random initializations of the same model configuration. Task-models are sorted by performance on V4. Hues represent cluster types (inset). The baseline is the average performance of an untrained network. **d,e**, Comparisons between pairs of task-clusters. Individual task-models as circles, and box-plots depict their distribution for each task-cluster. The number of stars represent the *p-value* upper thresholds (0.05, 10*^−^*^2^, 10*^−^*^3^, 10*^−^*^3^) of each comparison test between pairs of clusters. For each pair, we ran a pairwise Wilcoxon signed rank test between the group contrast performance of individual units *n* = 342 (V1), and *n* = 202 (V4), and applied Holm-Bonferroni correction to account for the six multiple comparisons. **f**, The normalized performance (increment over untrained baseline divided by the mean) for all individual tasks on V1 and V4. The variance across tasks in V4 is significantly different than in V1, rendering it more functionally specialized (*P* = .00135, *n* = 23, Levene’s test for equality of variances).

After determining the optimal layer, scale, and regularization parameters for each brain area and task network on a validation set, we found that V1 responses were better predicted than V4’s (Fig. 2c). The top *taskonomy* -based model performances (test correlation to average over trails [*±* standard error of the mean across seeds]) were 0.690 *±* 0.0003 (V1) and 0.412 *±* 0.0014 (V4). Equivalent results in terms of fraction of explainable variance explained (Methods, [20]) were 0.496 *±* 0.0005 (V1) and 0.125 *±* 0.0016 (V4) (Fig. S5). This performance discrepancy between V1 and V4 could be explained only partly by differences in selectivity between V1 and V4 responses (mean *±* s.d. selectivity indices of V1: 0.40 *±* 0.23, V4: 0.56 *±* 0.26; two-sided *t*-test: *t*(542) = *−*6.728*, p* = 4 *·* 10*^−^*^11^; Fig. S2b–c), because the selectivity index of individual neurons was negatively correlated with the performance yielded by our top model (*ρ* = *−*0.35*, p <* 10*^−^*^8^ on V4 neurons; Fig. S2e).

Pre-training was very effective: Most models widely outperformed the untrained model baseline with random weights – unlike earlier work in the mouse visual cortex [24]. The two exceptions on V4 were texture edges and 2D keypoints (Fig. 2c), which did not improve over the untrained baseline. These results are in line with previous work [25] where models using pretrained representations significantly outperformed untrained representations fitted to human fMRI responses in inferior temporal cortex (IT).

### Semantic classification tasks predict V4 best, while V1 is well-predicted by diverse tasks

The best predictive task-models on V4 were the two semantic classification tasks: scene classification (0.4117 *±* 0.0013), and – consistent with prior work [8, 9] – object classification (0.4089 *±* 0.0010). In contrast, we found that in V1 the top models with comparable performance were diverse and not specifically tied to semantic-related tasks – they came from semantic, 3D, and 2D task groups: semantic segmentation (0.6900 *±* 0.0003), 2D segmentation (0.6886 *±* 0.0010), euclidean depth (0.6898 *±* 0.0007) (Fig. 2c, left). Interestingly, the performance of semantic segmentation in V4 (a pixel-to-pixel task), did not yield a high performance in comparison (0.3739 *±* 0.0008), while it was among the best-performing tasks on V1 (see Discussion).

To unveil trends that apply beyond individual tasks, but that are consistent for functionally related task-groups (i.e. semantic, geometric, 3D, and 2D task clusters), we compared the average performances of each task cluster. Consistent with our previous results at the individual task level, we found evidence at this coarser level for the specialized role of V4 towards semantic tasks: there were significant differences between all pairs of clusters with the semantic group on top. In contrast to observations at the individual task level, and in line with previous work [20, 26], we found that semantic representations were significantly better at predicting V1 responses than other groups (Fig. 2d) and identified significant differences between all pairs of groups (except between 2D and geometric). Nevertheless, when looking at the group median performances, we found that the group type was less critical in V1 than V4: all groups in V1 were above 74% of the gap between the untrained baseline and top median group (semantic), while the lowest group median in V4 (2D) reached only 16% of that gap (Fig. 2d, e).

### V4 is more specialized than V1

How important is the specific task objective over the untrained baseline to obtain better predictive performances? We found that the choice of task (and task cluster as shown before) did not affect performance as much in V1 as it did in V4 (Fig. 2d–f). We quantified this functional specialization by computing the variance of the performance of all task-models normalized to their mean (excluding the untrained network). The variance for V4 was higher than for V1 (*p <* 0.01, *n* = 23, Bartlett’s test after Fisher’s *z*-transformation; Fig. 2f), rendering it more specialized. These results back the traditional notion of generality in the features extracted by V1, as they support multiple downstream tasks, while they also highlight the more specialized role of V4 in visual processing.

### Building models that jointly read out from pairs of tasks

Our comparisons at the individual and cluster levels suggest that semantic tasks drive representations that best match ventral visual areas in the brain, especially area V4. However, semantic tasks outperformed the next-best predictive tasks only by a relatively small margin (Fig. 2c, right), making it difficult to dismiss these other tasks altogether as computational goals of V4 function.

Moreover, the task-models that directly followed object classification – reaching 81% of the gap between the untrained and scene classification model – extract features relevant for 3D understanding (3D keypoints, 2.5D segmentation, and surface normals estimation) which could be aligned with V4 functions – as recent studies suggest that many cells in V4 are tuned to solid-shape (3D) properties [16]. For example, the 3D keypoints task aims to find points of interest that could be reliably detected even if an object in the scene is observed from different perspectives, and then to extract local surface features at these points. Detecting 3D borders is essential to solve this task because these keypoints tend to be around object corners [27], likely capturing useful information for downstream invariant object recognition.

Do computational goals (like these 3D-related objectives) provide representations that explain aspects of V4 computation beyond those explained by semantic tasks alone? We addressed this question by comparing the individual task-model performances to the performance of a single model that jointly reads out from pairs of pretrained feature spaces (Fig. 3a). If we find that a larger response variance is explained by adding a second set of features extracted by a different task-network to the first one, we could claim that the nonlinearities of second network are likely not implemented by the first one. Otherwise, the original readout of the single-task model could have exploited them to yield a higher performance.

**Figure 3.**
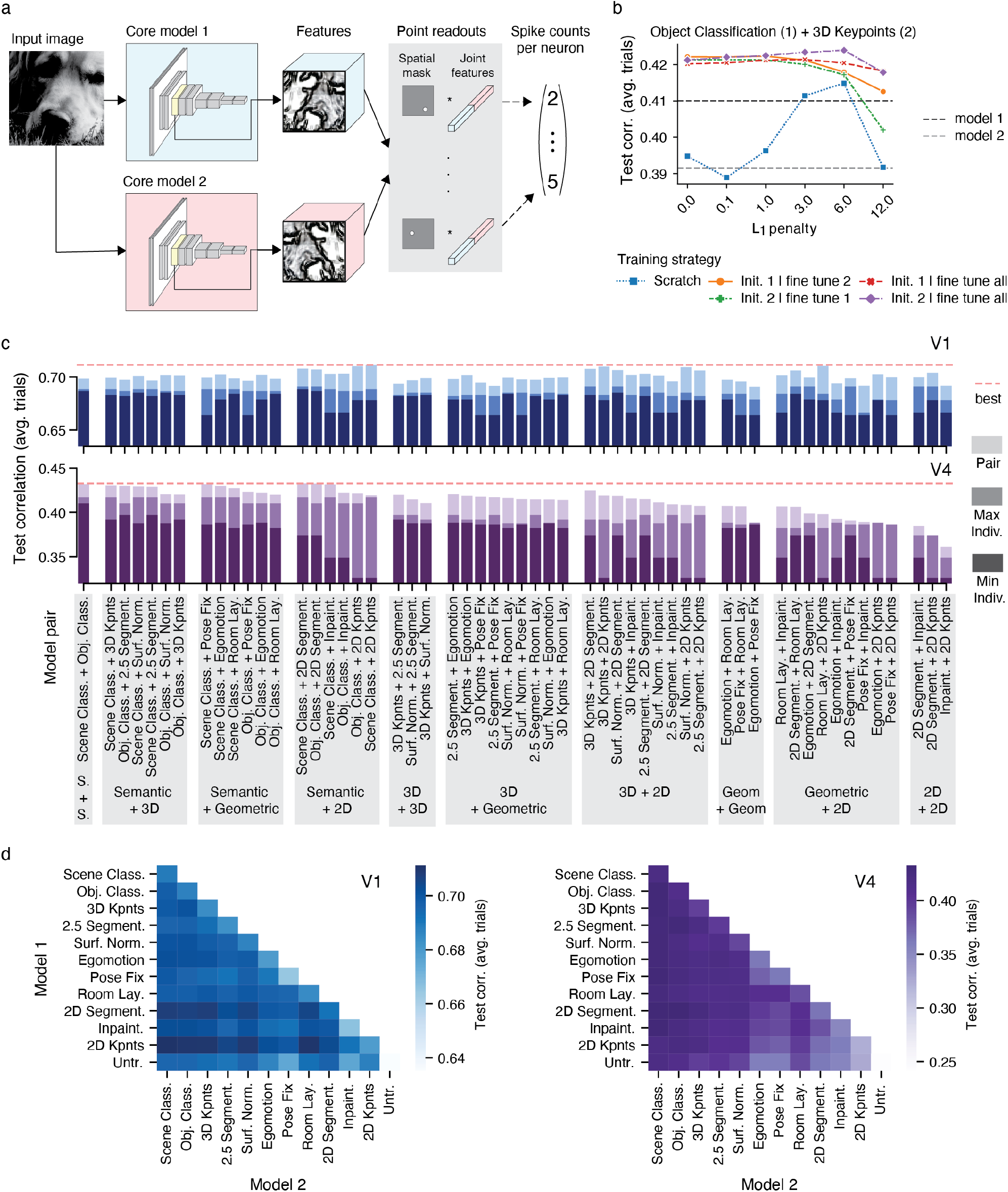
Jointly reading from pairs of tasks. **a**, Modeling approach. We modified the methods described in Fig. 1 to simultaneously read out neural responses from two feature spaces. We forwarded the same input image, at the same scale, to two pretrained *taskonomy* networks, extracted the features at the same layer and concatenated them. We then learned a point readout [17] for each neuron and always keep the cores’ weights frozen during training. **b**, Comparison of training strategies when jointly reading from object classification (core 1) and 3D keypoints (core 2). The resulting increased dimensionality makes it harder to find global optima, so we compared – under several regularization strengths (*L*_1_ penalty) –five strategies that leverage the individual task-models’ optimum readout weights as per validation set from the models we trained earlier. We trained from scratch (i.e. initialized the readout weights at random), initialized the readout of core 1 (2) and finetuned the readout of core 2 (1), and initialized the readout of core 1 (2) and finetuned all readout weights. The individual task-model performances are in dotted lines for comparison. **c** Comparison of pair task models in V1 (top) and V4 (bottom). Pairs are grouped based on the task-group identities of pair members. In dark (middle-dark) color saturation, we show the worse (best) individual task model from the pair. In lightest color saturation, we show the performance of the pair model. The highest performing pair model is shown as a red dotted line to facilitate comparisons. Performances are shown in terms of correlation between model predictions and average over trials. Baseline on all bar plots is the performance of the untrained network. **d** Heat map representation of pair task models’ performances in V1 (let) and V4 (right). Here, in addition to the pairs in **c**, the pair models with identical members (diagonal), and pairs built between tasks and the untrained network (bottom row) are included.

In practice, however, it is generally harder to find global optima with an increased set of input features when the amount of training data is kept the same. Naively training the joint readout of a pair of tasks from scratch could lead to worse performance than the individual task-model performances. Therefore, we validated different training strategies together with an *L*_1_ regularization penalty (Fig. 3 b): (1) The readout weights were initialized randomly, and we learned them from scratch, (2) The readout weights corresponding to the first core were initialized with the optimal ones found previously (as per validation set), while those corresponding to the second core were initialized with zeroes. Then we fine-tuned the readout weights of the second core only. This is approach was repeated by swapping the order of the cores. (3) We repeated the second approach, but instead of fine-tuning only the second core readout weights, we fine-tuned all.

We built and trained models for pairs of tasks following these strategies. In addition to scene and object classification, we considered the top three tasks on V4 of each group (Fig. 2c, right), and the untrained baseline; and built models with all possible pairwise combinations (Fig. 3c–d). This means 12 individual tasks and 78 pair task-models per brain area (Fig. 3d).

We observed that initializing a portion of the readouts with the already optimized ones consistently outperformed training from scratch, as illustrated in an example involving object classification and 3D keypoints (Fig. 3b). Moreover, in most cases, fine-tuning all readout weights yielded better results than updating only some of them.

### Non-semantic tasks contribute useful nonlinearities beyond those provided by individual semantic classification tasks

We found that the performance of pair task-models was always better than any of their individual constituents (Fig. 3c). The average performance improvements over the highest-performing individual task in each pair were 3.9% (V4) and 2.2% (V1). The largest increase in performance in V4 came from a pair comprising 3D keypoints and 2D segmentation, which resulted in a 8.5% improvement over 3D keypoints alone. Similarly, the largest boost in V1 was obtained from a pair consisting of 2D keypoints and 3D keypoints, leading to a 4.0% increase over 2D keypoints alone.

We conducted two control experiments. First, to determine whether the observed performance gains may simply be due to the additional nonlinear capacity of the models, we trained a set of pair task-models by pairing each individual task with an untrained model. In every instance, the performance was found to be superior when pairing with another task instead of the untrained baseline (Fig. 3d; last row). Second, training models with higher-dimensional feature spaces required adapting the hyperparameters of the non-convex optimization procedure (regularization, number of iterations, etc.). To ensure that these changes did not trivially lead to a better model, we trained pair task-models consisting of one task’s features simply duplicated (Fig. 3d; diagonals). We found that incorporating the nonlinearities of a different task always resulted in better performance than a pair of duplicate task features (Fig. 3d). These results suggest that the improved performance observed in V1 and V4 is attributed to the additional nonlinear computations provided by the supplementary tasks rather than just the enhanced flexibility resulting from a greater feature dimensionality.

### Semantic features are critical to explain the largest fraction of variance in V4

We found that 2D tasks – in particular 2D keypoints – were consistently among the highest performing pairs in V1: 2D keypoints + scene classification (0.711), room layout + 2D keypoints (0.710), object classification + 2D keypoints (0.710), 3D keypoints + 2D keypoints (0.709), surface normals + 2D keypoints (0.709), scene classification + 2D segmentation (0.708). Interestingly, the individual 2D keypoints model was not among the top ten tasks in V1, but pairing it with members of other task groups yields the best performing models, suggesting that 2D keypoints is the most non-redundant task with other top-performing V1 tasks.

On the other hand, we found that the semantic classification tasks were consistently among the top pairs in V4 (Fig. 3 c): scene classification + object classification (0.432), scene classification + 2D segmentation (0.433), scene classification + fixed camera pose (0.432), scene classification + inpainting (0.431), object classification + 2D segmentation (0.432), scene classification + 3D keypoints (0.431).

To investigate whether the nonlinearities captured by scene and object classification are equivalent to those captured by other pairs of tasks exhibiting similar performance, we extended our previous methods (Fig. 3a) to jointly read from triplets of tasks.

Following our previous approach, we fine-tuned all readout weights after initializing the readouts corresponding to two task cores with the optimal weights obtained from pair task models (Fig. 3) and the weights corresponding to the third core with zeroes. We focused on studying improvements provided by the features of a third task core over the object + scene classification pair. We observed only minimal improvements over this pair (0.4 *±* 0.1%) ranging from 0 to 1.1%. Controls involving an untrained, scene, and object classification tasks as a third core yielded no improvements. These improvements are within the variability range obtained across seeds and are much smaller than than improvements provided by pairs of tasks over individual ones. Therefore, most nonlinearities captured by non-semantic tasks that are helpful to predict V4 responses can emerge from purely semantic training objectives.

### Data-rich and robust models predict V1 and V4 better

Taskonomy networks helped us addressed a prevalent issue in most goal-driven system identification studies where multiple sources of variability across models are simultaneously at play (e.g. architecture, training dataset, training strategies, computational goal). By sharing architecture, training dataset, and optimization methods, these networks facilitated a fair comparison of the computational objective. To make progress in understanding the effects of other sources of variation, we built V1 and V4 models that use the representations of widely used, state-of-the-art networks that were largely trained on ImageNet [21] and had varying architectures and different training strategies (Fig. 4): AlexNet [32], VGG19 [31], Cornet-S [33], Resnet50 [18], SimCLR [28], Resent50 with adversarial training (Robust *L*_2_*, ɛ* = 0.1) [30], and Resnet50 trained on ImageNet and Stylized ImageNet (StyleImNet) [29].

**Figure 4.**
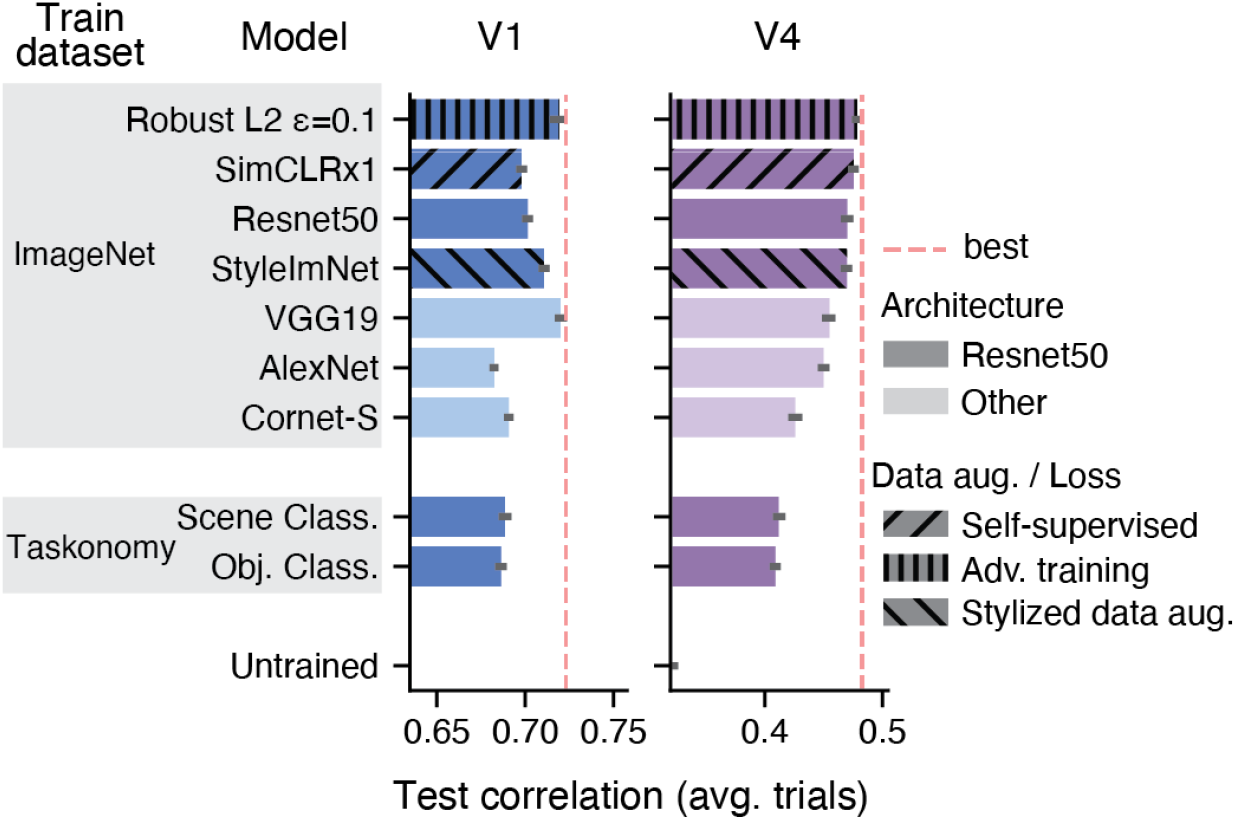
Comparisons with ImageNet-trained networks. We considered CNNs trained on the ImageNet dataset with varying architectures, data augmentation strategies, and losses (supervised vs. self-supervised) and compared them with the *taskonomy* semantic classification networks on V1 (left) and V4 (right). Red dotted line indicates the performance of the best model on each visual area. We found that ImageNet networks were better than *taskonomy* on V4 for any architecture. Self-supervised (SimCLRx1, [28]) and shape-biased (StyleImNet Resnet50, [29]) networks yielded comparable performance in V4 to the supervised, texture-biased Resnet50. The highest performing network on V4 was the adversarially robust Resnet50 [30]. In V1, our top *taskonomy* networks were comparable to AlexNet and Cornet-S but performed worse than the ImageNet counterpart with matching architecture (Resnet50). VGG19 [31] and the robust Resnet50 were the best match to V1.

We found that – beyond the semantic computational objective – the training dataset is critical in driving representations that best match V4 responses: all ImageNet models yielded higher performance than the top semantic *taskonomy* networks (Fig. 4; right). Our results also suggests that a Resnet50 architecture is generally better than others (VGG19, AlexNet, Cornet-S). Interestingly, a shape-biased network (StyleImNet) does not lead to better predictions than a texture-biased counterpart (Resnet50), even though, at the behavioural level, the visual system exhibits bias to shapes [29]. Furthermore, in line with previous work [34], we found that self-supervised representations from SimCLR had comparable performance to the supervised-trained Resnet50. Although not explicitly trained on object classification, SimCLR provides useful features for this task: a shallow multi-linear-perceptron readout with little supervision achieves competitive accuracy on ImageNet [28]. Finally, we obtained the highest V4 performance (0.4786 *±* 0.0011) with the adversarially robust Resnet50 that was trained to produce stable classification outputs under small perturbations of at most 0.1 *L*_2_ size. We observed that increasing the size of this robustness ball led to poorer performance in V4 – as it also negatively affects ImageNet top1 accuracy [30]. Overall, these results indicate that the match to V4 neural data can be strongly influenced by the training dataset, architecture, and adversarial robustness.

Our V1 results revealed trends that partially differ from those observed in V4. First, we found that some ImageNet-trained models (AlexNet and Cornet-S) yielded comparable performance to the top semantic *taskonomy* -based models. However, when using the same Resnet50 architecture, we obtained better performance with ImageNet models. As in V4, self-supervised learning and shape-biased networks performed similarly to their supervised, texture-biased counterpart. However, our best V1 models came from VGG19 (0.7199 *±* 0.0008) – in line with [20] –, and the robust Resnet50 (0.7184 *±* 0.0027), which is consistent with findings reported in [26] and [35]. In a similar way to V4, increasing the robustness size of the ResNet50 model led to poorer performance. Overall, our results suggest that data-rich and robust models are more effective in capturing V1 and V4 representations.

## Discussion

A common challenge when comparing models in task-driven system identification studies is that multiple axes of variability get confounded, making it difficult to draw conclusions about individual factors like architecture or training objectives. We explored multiple normative accounts beyond object classification of V4 and V1 single neuron responses to natural stimuli using the representations of *taskonomy* networks as they have matching architectures and were trained on the same dataset but different computer vision tasks. Beyond solidifying existing evidence suggesting that semantic goals (e.g. object classification) drive representations that best match ventral stream responses [8, 20, 22, 36, 37], our results revealed a high task specialization of V4 function towards semantic tasks in contrast to the more general representations in V1 that can support multiple downstream tasks. Moreover, when predicting V4 responses from pairs of different tasks features, we found that non-semantic tasks contribute useful nonlinearities over individual semantic ones. However, a model that jointly reads from a pair of semantic-driven features alone was among the best of our *taskonomy* -based models of V4, suggesting that semantic objectives can sufficiently capture the nonlinearities explained by other models.

Over the last decades, the functional role of V4 has been characterized by identifying neuronal tuning directions using creatively synthesized, parametric stimuli (see [1] for a review). This powerful approach has led to multiple insights about V4’s preferences, but does require generating hypotheses of stimulus features (e.g. tuning for blurry vs. hard edges) and painstaking experiments to test each of them. Based on the patterns of predictive performance across tasks (and task-pairs) our systems identification approach provides another lens on how we can better understand V4 function. Importantly, our results suggest that these two approaches can inform each other. Specifically, our work predicts a strong relationship between 3D visual processing and V4 function, shifting the long-standing focus of V4’s functional role in flat shape processing: 3D representations ranked high after semantic ones (Fig. 2), and they captured novel nonlinearities over those of individual semantic tasks (3). This supports recent discoveries showing that some V4 cells encode solid shape (3D) features [16]. A promising future research direction is to explore the relationship between normative accounts of V4 at the population level and individual cell properties, specially now that we can first probe multiple stimulus features *in-silico* using our best CNNs before moving to the expensive *in-vivo* alternatives.

Our results revealed the affinity of V1 to 2D processing, consistent with classical views of V1 function [38]. We found that 2D segmentation was among the top *taskonomy* networks (Fig. 2), while it also maintained high predictive performance on the deepest layers that explicitly solve the task Fig. S3. Moreover, we found that the 2D keypoints tasks was consistently among the highest performing pairs (Fig. 3). This task approximates the output of a classical computer vision algorithm (SURF [39]) that was engineered to identify points of interests in an image (using an approximation of multi-scale difference-of-Gaussians [DoG]) and to extract features around them (e.g. orientation tuning) that facilitate matching these points across affine transformations of the image (rotation, translation, scaling, shear mapping). This sequence of operations (multi-scale DoGs, and identifying orientation preference with rotated wavelets) bears similarities with classical descriptions of V1 function [38]. Finally, we found that ImageNet-trained CNNs (Fig. 4) yielded our best performing V1 models. This observation is consistent with recent observations [20, 26] claiming that object classification goals induce useful V1 nonlinearities beyond energy models using multi-scale Gabor features [40], and that these CNNs can learn known nonlinear phenomena like cross-orientation inhibition [41] that those models miss [42].

We found that in contrast to object and scene classification tasks, semantic segmentation was not as effective at predicting V4 responses, while it was among the best in V1 (Fig. 2). One likely explanation has to do with the architecture of the *taskonomy* semantic segmentation network: unlike modern networks like U-Net [43] or Mask R-CNN (with an FPN backbone [44]) [45] that include lateral connections between encoding and decoding streams at different scales, the decoder of the *taskonomy* network has only access to the high-level output of the encoder to infer a segmentation mask. To keep the architecture of the core consistent across tasks, we considered the encoder’s representations. While the early encoder layers capture rich enough representations to predict V1, the later layers underperform on V4, possibly because 1) they miss task-relevant nonlinearities that are only implemented by the decoder network, and 2) they must maintain details about object boundaries at the pixel level in their representations due to the lack of multi-scale lateral connections provided to the decoder. This is particularly relevant because it contrasts with V4 units, which have been shown to exhibit a degree of translation invariance [9, 46].

We found that V1 was better predicted by task representations than V4 (Fig. 2, Fig. S5), even though the stimulus-driven variability was comparable between these two areas. We do not believe that the seemingly low explained variance by our V4 models lessens the conclusions drawn in this paper: How well a model performs in absolute terms depends on many experimental factors including the amount of data available to fit the model, the signal-to-noise ratio of the model, the nature of the pre-training dataset etc. In our work, we believe that the reasons for the comparably low values in V4 are two-fold. First, these results are likely due to our experimental design that maintained the sequence of images (without interleaving blanks) on the repeated test-trials. Although this choice makes history effects in V4 visible so future (dynamic) models can account for them, it prevents averaging out the variability introduced by the previous image, because the static image-based models cannot account for it. The dynamic effects are likely stronger in V4 than in V1 because the latter – an earlier visual area – has lower response latency and the image display times (120 ms) may be sufficient to reach stable dynamics (see Fig 2b from [20]). In contrast, display times of 120 ms in V4 have been used to model “core object recognition” – a term coined for the first feed-forward wave of visual processing [8]. Second, the models we compare were trained on the *taskonomy* dataset, whose image statistics deviate from those of the ImageNet images shown in the experiments. Hence, it is expected that these models do not perform on par with a model that was trained in-domain. We do not think that this weakens our conclusions because all *taskonomy* models have the same systematic disadvantage of having to generalize somewhat beyond their training set in terms of image statistics. Because we do not have diverse task labels for ImageNet, we make the assumption that the variable of interest (training objective) does not interact with the image statistics. To verify that the lower performance of our models is indeed caused by those two factors discussed above and not related to highly suboptimal modeling (e.g. readouts, hyperparameter optimization, etc.), we conducted additional control experiments. When using the same data-rich cores as current state-of-the-art models, we found that our models perform similarly to those displayed on the BrainScore leaderboard [36]. We used ImageNet-trained models (Fig. 4) and found that our best V4 model was a Resnet50 trained to be robust to small adversarial perturbations [30]. The performance of this model in V4 in terms of correlation coefficient was *∼* 0.48, which is in the same ballpark as the top models on the Sanghavi, et al. (2021) [47] benchmark on BrainScore to date.

We found significant differences between task groups (Fig. 2 d–e), but the absolute value of performance differences across tasks are seemingly small (test correlation values range from 0.3 to 0.48 in V4), which could potentially raise concerns about our claims. We think these concerns are unfounded for two reasons. First, we found a structured pattern of tasks performances that is largely consistent with expectations based on existing work and far from random. As discussed before, our results predict V4’s affinity to semantic and 3D tasks, and highlight the important role of semantic and 2D representations in V1. Second, we argue that small differences in test performance may carry meaningful implications about the ability of models to capture neural phenomena. For example, in a previous study [42], we found that the difference between an energy model using a Gabor filter bank (GFB) and a one-layer CNN with a divisive normalization nonlinearity (CNN+DN) was seemingly small (*∼* 0.05 of FEVE) in favour of the latter. However, when running *in-silico* experiments, it was found that CNN+DN was able to capture cross-orientation inhibition – a known nonlinear phenomenon in V1

– while the GFB model was not. Thus, the difference we find between task performances
– although small – can be consequential and should not be quickly dismissed. A promising future direction is to leverage *in-silico* experiments to find stimuli that bring to light the differences between models, for example via controversial stimuli [48] that drive one model’s predictions while not the other(s).

In contrast to similar efforts applied to measurements of aggregated neural activity (i.e. fMRI data) from multiple visual areas [12, 49], we evaluated the *intermediate* representations of *taskonomy* networks – not just their final layers – to evaluate their affinities to single-cell responses. We indeed found that intermediate layers provided the best match to the data – regardless of the task – (Fig. S6, Fig. S4) and that diverse tasks offered comparable top performances in V1 and mostly semantic tasks were optimal in V4. [12] considered large, separate regions of interest (OPA, PPA, LOC, EarlyVis, and RSC) in human fMRI and found that object and scene classification were best across areas and subjects. V1 and V4 would likely fall into the same “EarlyVis” ROI, preventing separate conclusions of each of these areas. Dwivedi et al. (2021) [49] did considered more granular areas of the ventral stream including V1 and V4, and found that 2D tasks best explain V1 (consistent with our results) and that – in contrast to our findings – 2D tasks best explain V4. This could be attributed the nature of fMRI where the nonlinear properties of individual cells can be averaged out, and last-layer 2D representations are more linear than other tasks (see similarity across layers in [15]). Overall, our work at the individual cell level distill more insights about V1 and V4 function than these other data modalities.

Although we account for architecture, training dataset, and optimization methods to be equivalent across core representations, a potential concern not addressed in our study is that there is evidence for variability in intermediate representations of identical networks trained on the same task but initialized differently [50]. This is a challenge faced by most goal-driven system identification studies and it is constrained by the unavailability of multiple instances of a task-network – as it is the case for *taskonomy* encoder networks. That said, we remark that the representational consistency across instances is high for early to mid layers, decreasing with network depth [50]. V1 and V4 were best explained by early to mid levels. Thus, the effects of initialization are not expected to be as strong as if later layers were used. For example, in the concrete case of AlexNet [32], an ImageNet-trained network considered in our work and in the study by [50], we found that the best predictive layer of V4 was *Layer 3*, which had a representational consistency above 0.95 (on a scale of 0–1), similar to that of the earliest layers (see Fig. 6 from [50]). Moreover, we did consider multiple tasks for each of the task groups (semantic, 3D, geometric, and 2D) and found significant differences at the group level (Fig. 2, Fig. S7) that support our main claim favouring the semantic specialization of V4. We make our data and code available to facilitate further progress identifying the effects of network initialization.

Are semantic training objectives critical to drive good representations of V1 and V4? We found that a self-supervised network (SimCLR) worked just as well than Resnet50 in line with [34]. Although these self-supervised methods contain no explicit semantic training objective, their inherent data augmentation strategies enforce similar invariances as required for object recognition [51] and their learned representations predict semantic labels well [28]. Moreover, we found that additional data augmentations, particularly those provided by adversarial training, were very effective at improving our models’ performance in both areas. These results are aligned with what people have found in V1 [26, 35] that suggest that robustness to perturbations explain V1 responses better. Similar work that goes in the other direction, has shown that making a CNN more brain-like by co-training on object classification and to predict neural responses, makes CNNs more robust to small perturbations [52].

Taken together, our results provide evidence for semantic tasks as a normative account of area V4 and predict that particular tasks (e.g. 3D) have strong affinities to V4 representations. Although we are yet to explain most variance, our findings suggest that promising directions to improve the predictive power of V4 responses require using data-rich, robust models (also shown to be useful to match behavioural phenomena [53]), and architectural changes (e.g. recurrence, top-down processing streams) that account for dynamical processing in the brain [54]. Moreover, our multi-task modeling approach, together with *in-silico* experimentation, promises to facilitate the generation of hypotheses about tuning directions via maximally exciting inputs (MEIs) [10, 11, 55], diverse exciting inputs (DEIs) [56, 57], or controversial stimuli [48].

## Materials and methods

### Ethics statement

All behavioral and electrophysiological data were obtained from two healthy, male rhesus macaque (*Macaca mulatta*) monkeys aged 15 and 16 years and weighing 16.4 and 9.5 kg, respectively, during the time of study. All experimental procedures complied with guidelines of the NIH and were approved by the Baylor College of Medicine Institutional Animal Care and Use Committee (permit number: AN-4367). Animals were housed individually in a large room located adjacent to the training facility, along with around ten other monkeys permitting rich visual, olfactory and auditory interactions, on a 12h light/dark cycle. Regular veterinary care and monitoring, balanced nutrition and environmental enrichment were provided by the Center for Comparative Medicine of Baylor College of Medicine. Surgical procedures on monkeys were conducted under general anesthesia following standard aseptic techniques. To ameliorate pain after surgery, analgesics were given for seven days.

### Electrophysiological recordings

We performed non-chronic recordings using a 32-channel linear silicon probe (NeuroNexus V1x32-Edge-10mm-60-177). The surgical methods and recording protocol were described previously [19]. Briefly, form-specific titanium recording chambers and headposts were implanted under full anesthesia and aseptic conditions. The bone was originally left intact and only prior to recordings, small trephinations (2 mm) were made over medial primary visual cortex at eccentricities ranging from 1.4 to 3.0 degrees of visual angle. Recordings were done within two weeks of each trephination. Probes were lowered using a Narishige Microdrive (MO-97) and a guide tube to penetrate the dura. Care was taken to lower the probe slowly, not to penetrate the cortex with the guide tube and to minimize tissue compression).

### Data acquisition and spike sorting

Electrophysiological data were collected continuously as broadband signal (0.5Hz–16kHz) digitized at 24 bits. Our spike sorting methods mirror those in [19, 20], code available at https://github.com/aecker/moksm. We split the linear array of 32 channels into 14 groups of 6 adjacent channels (with a stride of two), which we treated as virtual electrodes for spike detection and sorting. Spikes were detected when channel signals crossed a threshold of five times the standard deviation of the noise. After spike alignment, we extracted the first three principal components of each channel, resulting in an 18-dimensional feature space used for spike sorting. We fitted a Kalman filter mixture model to track waveform drift typical for non-chronic recordings [58, 59]. The shape of each cluster was modeled with a multivariate t-distribution (*df* = 5) with a ridge-regularized covariance matrix. The number of clusters was determined based on a penalized average likelihood with a constant cost per additional cluster [60].

Subsequently, we used a custom graphical user interface to manually verify single-unit isolation by assessing the stability of the units (based on drifts and health of the cells throughout the session), identifying a refractory period, and inspecting the scatter plots of the pairs of channel principal components.

### Visual stimulation and eye tracking

Visual stimuli were rendered by a dedicated graphics workstation and displayed on a 16:9 HD widescreen LCD monitor (23.8”) with a refresh rate of 100 Hz at a resolution of 1920 *×* 1080 pixels and a viewing distance of 100 cm (resulting in *∼* 63*px/^◦^*). The monitors were gamma-corrected to have a linear luminance response profile. A camera-based, custom-built eye tracking system verified that monkeys maintained fixation within *∼* 0.95*^◦^* around a *∼* 0.15*^◦^*-sized red fixation target. Offline analysis showed that monkeys typically fixated much more accurately. After monkeys maintained fixation for 300 ms, a visual stimulus appeared. If the monkeys fixated throughout the entire stimulus period, they received a drop of juice at the end of the trial.

### Receptive field mapping and stimulus placing

We mapped receptive fields relative to a fixation target at the beginning of each session with a sparse random dot stimulus. A single dot of size 0.12*^◦^*of visual angle was presented on a uniform gray background, changing location and color (black or white) randomly every 30 ms. Each fixation trial lasted for two seconds. We obtained multi-unit receptive field profiles for every channel using reverse correlation. We then estimated the population receptive field location by fitting a 2D Gaussian to the spike-triggered average across channels at the time lag that maximizes the signal-to-noise-ratio. During V1 recordings, we kept the fixation spot at the center of the screen and centered our natural image stimulus at the mean of our fit on the screen (Fig. 1b). During V4 recordings, the natural image stimulus covered the entire screen. We accommodated the fixation spot so that the mean of the population receptive field was as close to the middle of the screen as possible. Due to the location of our recording sites in both monkeys, this equated to locating the fixation spot close to the upper border of the screen, shifted to the left (Fig. 1c).

### Natural image stimuli

We sampled a set of 24075 images from 964 categories (*∼* 25 images per category) from ImageNet [61], converted them to gray-scale (to be consistent with similar system identification studies and reduce complexity), and cropped them to keep the central 420 *×* 420px. All images had 8 bit intensity resolution (values in [0,255]). We then sampled 75 as our *test-set*. From the remaining 24000 images, we sampled 20% as *validation-set*, leaving 19200 as *train-set*. We used the same sets of images for V1 and V4 recordings. During a recording session, we recorded *∼* 1000 successful trials, each consisting of uninterrupted fixation for 2.4 seconds including 300ms of gray screen (128 intensity) at the beginning and end of the trial, and 15 images shown consecutively for 120ms each with no blanks in between. Each trial contained either train and validation, or test images. We randomly interleaved trials throughout the session so that our test-set images were shown 40-50 times. The train and validation images were sampled without replacement throughout the session, so each train / validation image was effectively shown once or not at all. In V1 sessions, the images were shown at their original resolution and size covering 6.7*^◦^*(screen resolution of 63 pixels per visual angle). The rest of the screen was kept gray (128 intensity). In V4 sessions, the images were upscaled preserving their aspect ratio with bicubic interpolation to match the width of the screen (1920px). We cropped out the upper and bottom 420px bands to cover the entire screen. As a result, we effectively stimulated both the classical and beyond the classical receptive fields of both areas. Once the neurons were sorted, we counted the spikes associated to each image presentation in a specific time window following the image onset. These windows were 40-160ms (V1) and 70-160ms (V4).

### Explainable variance

A few isolated neurons were discarded if their stimulus-driven variability was too low [20]. The explainable variance in a dataset is smaller than the total variance because the observation noise prevents even a perfect model to account for all the variance in the data. Thus, targeting neurons that have sufficient explainable variance is necessary to train meaningful models of visually driven responses. For a neuron’s spike count *r*, the explainable variance Var_exp_[*r*] is the difference between the the total variance of all observed responses Var[*r*] and the variance of the observational noise *σ*^2^,

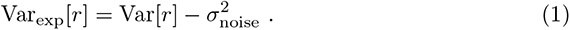

We estimated the variance of the observational noise by computing the variance of a neuron’s response *r_t_* in multiple trials *t* in which we presented the same stimulus *x_j_* and subsequently taking the expectation *E_j_*over all images,

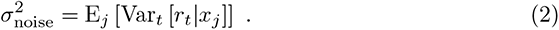

Neurons for which the ratio between the explainable to total variance (Eq. 3) was below 0.15 were removed. The resulting dataset includes spike count data for 202 (V1) and 342 (V4) isolated neurons, with an average ratio of explainable to total variance (s.d) of 0.306(0.181) and 0.323(0.187), respectively (Fig. S2). All variances were computed using the unbiased estimator and on the test-set responses due to the available repeated trial presentations.

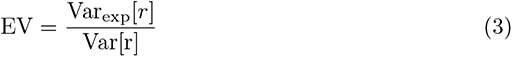

### Measuring Sparseness

We computed the selectivity index [62] (*SI*) as a measure of sparseness for every neuron on its test-set average responses. To do this, we first plotted the fraction of images whose responses where above a threshold, as a function of normalized thresholds. We considered 100 threshold bins ranging from the minimum to the maximum response values. The area under this curve (*A*) is close to zero for sparse neurons, and close to 0.5 for a uniform distribution of responses. Thus, following Quiroga et. al (2007) [62], we computed the selectivity index as *SI* = 1 *−* 2*A*. *SI* approaches 0 for a uniform distribution, and 1 the sparser the neuron is. In this study, we reported the mean and standard deviation of *SI* for both brain areas and found sparser responses in V4 than in V1 (see Results).

### Image preprocessing and resizing

An important step of our modeling pipeline was to adjust the size and resolution of the input images to our computational models (Supplementary Fig. S1). In V1, we effectively cropped the central 2.65*^◦^* (167px) at its original 63px*/^◦^* resolution and downsampled with bicubic interpolation to different target resolutions: 3.5, 7.0, 14, 21, 24.5, and 28px*/^◦^*. For practical and legacy reasons [20], in our codebase we first downsampled the images to a resolution of 35px*/^◦^*, followed by cropping and another downsampling step to obtained the target sizes and resolutions just reported. In V4, we cropped the images up to the bottom central 12*^◦^*, corresponding to 168px at the original 14px*/^◦^* resolution (63px*/^◦^ ×* 420*/*1920), in accordance with the neuron’s RF positions. These images were similarly downsampled to multiple target resolutions: 1.4, 2.8, 5.6, 8.4, and 11.2px*/^◦^*.

### Model architecture

Our models of cell responses consisted of two main parts: A pretrained core that outputs nonlinear features of input images, and a *spatial point readout* [17] that maps these features to each neuron’s responses. We built separate model instances for each visual area, input image resolution, task-dependent pretrained CNN, intermediate convolutional layer, regularization strength, and random initialization. Input images **x** were forwarded through all layers up to the chosen layer *l*, to output a tensor of feature maps *l*(**x**) *∈* R*^w×h×c^* (**w**idth, **h**eight, **c**hannels). Importantly, the parameters of the pretrained network were always kept fixed. We then applied batch-normalization [63] (Eq. 4), with trainable parameters for scale (*γ*) and shift (*β*), and running statistics mean (*µ*) and standard deviation (*σ*). These parameters were held fixed at test time (i.e. when evaluating our model). Lastly, we rectified the resulting tensor to obtain the final nonlinear feature space (Φ(**x**)) shared by all neurons, with same dimensions as *l*. The normalization of CNN features ensured that the activations of each feature map (channel) have zero mean and unit variance (before rectification), facilitating meaningfully regularized readout weights for all neurons with a single penalty – having input features with different variances would implicitly apply different penalties on their corresponding readout weights.

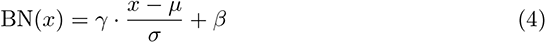

The goal of the readout was to find a linear-nonlinear mapping from Φ(**x**) to a single scalar firing rate for every neuron. Previous approaches have attempted to 1) do dimensionality reduction on this tensor and regress from this components (e.g. partial least squares) [8]; 2) learn a dense readout with multiple regularization penalties over space and features [20]; and 3) factorize the 3D readout weights into a lower-dimensional representation consisting of a spatial mask matrix and a vector of feature weights [64]. In this work we used the recently proposed spatial point readout [17] – also called *Gaussian readout* by the authors – that goes a step further and restricts the spatial mask to a single point. Per neuron, it computes a linear combination of the feature activations at a spatial position, parametrized as (*x, y*) relative coordinates (the middle of the feature map being (0, 0)). Training this readout poses the challenge of maintaining gradient flow when optimizing the objective function. In contrast to previous approaches that tackle this challenge by recreating multiple subsampled versions of the feature maps and learn a common relative location for all of them [65], the *Gaussian readout* learns the parameters of a 2D Gaussian distribution *N* (*µ_n_,* Σ*_n_*) and samples a location during each training step for every *n^th^* neuron. Σ*_n_* is initialized large enough to ensure gradient flow, and is then shrunken during training to have a more reliable estimate of the mean location *µ_n_*. At inference time (i.e. when evaluating our model), the readout is deterministic and uses position *µ_n_*. Although this framework allows for rotated and elongated Gaussian functions, we found that for our monkey data, an isotropic formulation of the covariance – parametrized by a single scalar *σ*^2^ – was sufficient (i.e. offer similar performance as the fully parametrized Gaussian). Thus, the total number of parameters per neuron of the readout were *c* + 4 (channels, bivariate mean, variance, and bias). Finally, the resulting dot product between the features of Φ(**x**) at the chosen location with an *L*_1_ regularized weight vector **w***_n_ ∈* R*^c^* was then followed by *f*, a point-wise nonlinear function ELU [66] offset by one (ELU + 1) to make responses positive (Eq. 5).

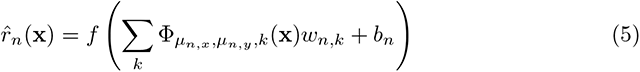

Beyond offering comparable performance compared to the factorized readout alternative with far less parameters, the most important motivation to use a single point readout was to make sure that all spatial nonlinear computations happen in the pretrained core feature extractor. We could draw mistaken claims about the nonlinear power of a feature space by computing new ones in the readout that combine rectified features computed at different spatial positions. For example, a readout that rectifies features produced by multiple simple cells with similar orientation at different locations can easily approximate phase invariance (i.e. complex cells) [38].

### Model training

We trained every model to minimize the summed Poisson loss across *N* neurons between observed spike counts *r* and our predicted spike rate *r̂* (Eq. 6, first term) in addition to the *L*_1_ regularization of the weights (Eq. 6, second term) with respect to the batch-normalization, and readout parameters.

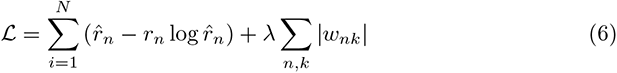

Since neurons across session from the same visual area didn’t necessarily *see* the same images (they were differently drawn over sessions), during each training step, we cycled through all sessions of the same visual area, sampling for each of them a fixed batch size of image-response pairs without replacement and kept track of the gradients of the loss with respect of our trainable parameters. Once a cycle was through, the gradients were added to execute an update of the weights of the weights based on the Adam optimizer [67] – an improved version of stochastic gradient descent. The initial learning rate was 3 *·* 10*^−^*^4^ and momentum 0.1. We continued to exhaust image-response pair batches from all sessions until the longest session was exhausted to count a full epoch. Once all image-response pairs had been drawn from a session, we restarted sampling batches from all available image-response pairs.

Every epoch, we temporarily switched our model into evaluation mode (i.e. we froze the batch-normalization running statistics), and computed the Poisson loss on the entire single trial *validation-set*. We then used early stopping to decide whether to decay the learning rate: we scaled the learning rate by a factor of 0.3 once the validation loss did not improve over five consecutive epochs. Before decaying the learning rate, we restored the weights to the best ones up to that point (in terms of validation loss). We ran the optimization until four early stopping steps were completed. On average, this resulted in *∼* 50 training epochs (or *∼* 40 minutes on one of our GPUs) per model instance.

### Taskonomy networks

The *taskonomy* networks [15] are encoder-decoder CNNs trained on multiple computer vision tasks. The original goal of the authors was to identify a taxonomy of tasks that would facilitate efficient transfer learning based on the encoder representations of these networks. Importantly for our study, all these networks were trained by the authors on the same set of images, which have labels for all tasks. These images consisted of 120k indoor room scenes. In this work, we used the encoder architecture of these networks, which was based on a slightly modified version of Resnet50 [18] that excluded average-pooling, and replaced the last stride 2 convolution with stride 1. However, these modifications did not change the number of output features of the intermediate layers we considered, keeping our *taskonomy* -based results fairly comparable with those of the original Resnet50.

The Resnet50 architecture – originally developed to solve ImageNet [61] – is made up of a series of hierarchical stages that include 1) an initial strided convolutional layer (conv1) followed by batch normalization, rectification, and max-pooling; 2) four processing layers, with 3,4,6, and 3 residual blocks, respectively; and 3) a final average pooling layer that maps features to the number of classes. Each residual block amounts to the rectified sum of two pathways: one that simply projects the input to the end (i.e. skip connection), and a second that consists of three successive convolutional layers with sizes 1, 3, and 1. In this work we trained models on the output of the first convolutional layer (conv1), and the output of the first residual block of each processing layer (i.e. layer1.0, layer2.0, layer3.0, layer4.0). The corresponding number of output feature maps (channels) for these layers was 64, 256, 512, 1024, and 2048, respectively.

We used several *taskonomy* encoder networks, listed in (Fig. 1c). The structure of the representations of these networks was presented by the authors via a metric of similarity across tasks: with agglomerative clustering of the tasks based on their transferring-out behavior, they built a hierarchical tree of tasks (see Figure 13 of their paper [15]). They found that the tasks can be grouped into 2D, 3D, low dimensional geometric, and semantic tasks based on how close (i.e. how similar) they are on the tree. We now briefly describe the tasks (for more details, see Supplementary Material from [15]):

#### 2D tasks

*Autoencoding* PCA finds a low-dimensional latent representation of the data. *Edge Detection* responds to changes in texture. It is the output of a Canny edge detector without nonmax suppression to enable differentiation. *Inpainting* reconstructs missing regions in an image. *Keypoint Detection(2D)* both detects locally important regions in an image (keypoints), and extracts descriptive features of them that are invariant across multiple images. The output of SURF [39] was the ground-truth output of this task. *Unsupervised 2D Segmentation* uses as ground-truth the output of Normalized cuts [68] which tries to segment images into perceptually similar groups.

#### 3D tasks

*Keypoint Detection (3D)* are like the 2D counterpart, but derived from 3D data, accounting for scene geometry. The output of the NARF algorithm [27] was the ground-truth output of this task. *Unsupervised 2.5D Segmentation* uses the same algorithm as 2D, but the labels are not only computed from RGB image, but also jointly from aligned depth, and surface normal images. It thus has access to ground-truth 3D information. *Surface Normal Estimation* are trained directly on the ground-truth surface normal vectors of the 3D meshes of the scene. *Curvature Estimation* extracts principal curvatures at each fix point of the mesh surface. *Edge Detection (3D)* (Occlusion Edges) are the edges where an object in the foreground obscures the background. It depends on 3D geometry and it is invariant to changes in color and lighting. In *Reshading*, the label for an RGB image is the shading function that results from having a single light point at the camera origin, multiplied by a constant albedo (amount of diffuse reflection of light radiation). *Depth Estimation, Z-Buffer*. *Depth Estimation, Euclidian* measures the distance between each pixel to the camera’s optic center.

#### Geometric tasks

*Relative Camera Pose Estimation, Non-Fixated* predicts the relative six degrees of freedom (yaw, pitch, roll, *x*, *y*, *z*) of the camera pose between two different views with same optical centers. *Relative Camera Pose Estimation, Fixated* is a simpler variant of the previous one where the center pixel of the two inputs is always the same physical 3D point – yielding only five degrees of freedom. *Relative Camera Pose Estimation, Triplets (Egomotion)* matches camera poses for input triplets with a fixed center point. *Room Layout Estimation* estimates and aligns 3D bounding boxes around parts of the scene. *Point Matching* learns useful local feature descriptors that facilitate matching scene points across images. *Content Prediction (Jigsaw)* unscrambles a permuted tiling of the image. *Vanishing Point Estimation* predicts the analytically computed vanishing points corresponding to an *x*, *y*, and *z* axis.

#### Semantic tasks

*Object Classification* uses knowledge distillation from a high-performing network trained on ImageNet [61] where its activations serve as a supervised signal (within the manually selected 100 object classes appearing in the *taskonomy* dataset). *Scene Classification* follows a similar approach, but uses a network trained on MITPlaces [69] with 63 applicable indoor workplace and home classes for supervised annotation of the dataset. *Semantic Segmentation* also follows the same supervised annotation procedure using a network trained on COCO [70] dataset with 17 applicable classes.

Finally, we included a control with matching architecture (Resnet50) but with random initialization.

### Other network architectures

In addition to the task-models based on *taskonomy* pretrained networks, we also built models with CNN feature extractors pretrained on the large image classification task ImageNet (ILSVRC2012) [61]. This is a dataset of 1.2 million images belonging to 1000 classes. In addition to the original Resnet50 [18] described before, we also considered other popular architectures (Supplementary Fig. S4): AlexNet [32], VGG19 [31], and Cornet-S [33].

AlexNet [32] consists of five convolutional layers with rectification, three max-pooling layers (between the first and second, second and third, and after the final convolutional layers), two fully connected layers after the last convolutional layer, and a final softmax layer. We used the output of all five convolutional layers in our study: conv1 1, conv2 1, conv3 1, conv4 1, conv5 1. Their number of output feature maps is 96, 256, 384, 384, and 256, respectively. We used the pretrained Pytorch implementation of this network found in the torchvision model zoo.

VGG19 [31] consists of 16 rectified convolutional layers that can be grouped into five groups (named conv1 to conv5) with 2, 2, 4, 4, and 4 convolutional layers with 64, 128 256, 512, and 512 feature maps, respectively; and a pooling layer after every group.

Finally, three fully connected and a softmax layers map the convolutional features to the 1000 predictions for each class. We used the original weights provided by [31], and not the default Pytorch version available in the torchvision model zoo.

Cornet-S [33] has a recurrent network architecture designed with known biological computations in the brain attributed to core object recognition. The network consist of a sequence of five main modules conveniently named V1, V2, V4, IT, and decoder. The V1 module consist of two subsequent convolutional layers with batch normalization and rectification with 64 channels in total; V2-IT are recurrent modules with an initial convolutional layer with batch normalization, and a recurrent series of three convolutional layers with rectification and batch normalization. The number of features are 128, 256, and 512 with 2, 4, and 2 recurrent time steps for V2, V4, and IT, respectively. The final decoder consist of average pooling, and a linear mapping to the 1000 output classes. The implementation of this network, including pretrained weights can be found here: https://github.com/dicarlolab/CORnet. We used the output of the first four modules of this network and found that V4 was best predicted by the corresponding V4 module, but our V1 data was best predicted (although only marginally better than V1) by the V2 module (Supplementary Fig. S4).

### Model configurations

In this study, we fitted a large set of task-models (*>* 10, 000) that include all viable combinations of 1) brain areas (2: V1, V4); 2) input resolutions (5); 3) pretrained CNNs (23 *taskonomy*, 1 random); 4) intermediate convolutional layers (5), 5) *L*_1_ regularization strengths (1-3 for most models); and 6) random initialization (5 seeds). Because of the receptive field size of higher layers in all networks, only large enough input resolutions were permitted in those cases. We used only 1-3 regularization penalties for most model configurations because we found that the optimal parameters from a fine-grained search of a single model where also appropriate for the corresponding layers of the *taskonomy* networks – actual optimal penalties led to negligibly differences in validation performance (within the noise of random seeds). The specific values we cross-validated over (after the fine-grained search) were: *λ*_conv1_ = *{*0.33, 1, 3*}, λ*_layer1.0_ = *{*3*}, λ*_layer2.0_ = *{*3, 6*}, λ*_layer3.0_ = *{*3, 9*}, λ*_layer4.0_ = *{*6, 12*}*.

### Performance evaluation

We computed the Pearson correlation between a model’s predictions with the average response over multiple presentations of our test-set to get comparable values to published results (e.g. [36]).

Furthermore, we also report the fraction of explainable variance explained (FEVE) (Fig. S5). The FEVE per neuron is given by Eq. 7

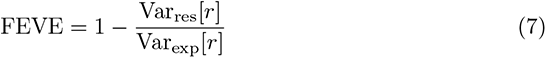

which utilizes the variance that is explainable in principle, Var_exp_[*r*] (Eq. 1), and the variance of the residuals corrected by the observation noise,

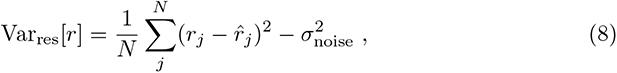

where *j* indexes images. This measure corrects for observation noise, which variance 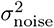 we estimated with Eq. 2. To compute model performance we averaged the FEVE across neurons.

## Data availability

The datasets of V1 and V4 recordings will be available for download at (https://figshare.com/s/30e05eb600646de94878) upon acceptance.

## Code availability

Our coding framework uses general tools like Pytorch [71], Numpy [72], scikit-image [73], matplotlib [74], seaborn [75], DataJoint [76], Jupyter [77], and Docker [78]. We also used the following custom libraries and code: neuralpredictors (https://github.com/sinzlab/neuralpredictors) for torch-based custom functions for model implementation, nnfabrik (https://github.com/sinzlab/nnfabrik) for automatic model training pipelines using DataJoint, nnvision (https://github.com/sinzlab/nnvision) for specific model definitions, ptrnets (https://github.com/sacadena/ptrnets) for readily available pretrained CNNs and access to their intermediate layers. Example code to train models on our data will be made available upon publication.

## Acknowledgements

The research was supported by the German Federal Ministry of Education and Research (BMBF) via the Competence Center for Machine Learning (FKZ 01IS18039A) and the Collaborative Research in Computational Neuroscience (CRCNS) (FKZ 01GQ2107 and NSF IIS-2113173); the German Research Foundation (DFG) grant EC 479/1-1 (A.S.E.), the Collaborative Research Center (SFB 1233, Robust Vision) and the Cluster of Excellence “Machine Learning – New Perspectives for Science” (EXC 2064/1, project number 390727645); the Bernstein Center for Computational Neuroscience (FKZ 01GQ1002); the National Eye Institute of the National Institutes of Health under Award Numbers R01EY026927 (A.S.T.), DP1 EY023176 (A.S.T.), and NIH-Pioneer award DP1-OD008301 (A.S.T), and the Intelligence Advanced Research Projects Activity (IARPA) via Department of Interior/Interior Business Center (DoI/IBC) contract number D16PC00003. This work was also supported by the National Institute of Mental Health under Award Number T32EY00252037. The U.S. Government is authorized to reproduce and distribute reprints for Governmental purposes notwithstanding any copyright annotation thereon. Disclaimer: The views and conclusions contained herein are those of the authors and should not be interpreted as necessarily representing the official policies or endorsements, either expressed or implied, of IARPA, DoI/IBC, or the U.S. Government. The funders had no role in study design, data collection and analysis, decision to publish, or preparation of the manuscript.

## Supplementary Figures

**Figure S1.**
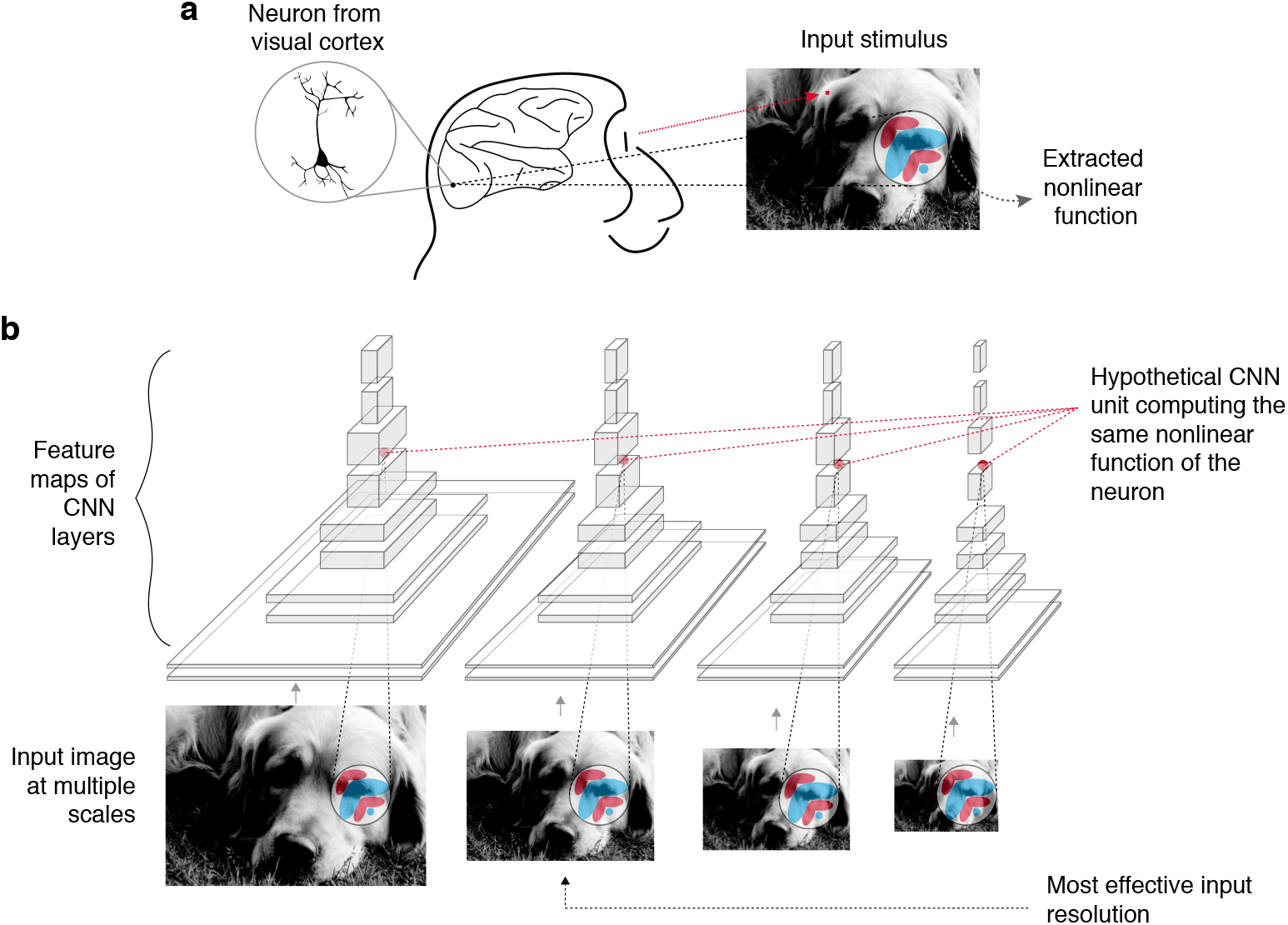
A case for input scale optimization. **a**, A single excitatory neuron from visual cortex, recorded from a head-anchored monkey sitting at a certain distance from a screen and fixating on a spot; extracts a nonlinear function of the input stimulus with a specific receptive field coverage. **b**, A pretrained deep convolutional neural network (CNN) extracts several nonlinear feature maps at each of its intermediate layers. A single output unit of a feature map computes a nonlinear function on its analytical receptive field with a fixed size in pixels. Even if the real neuron’s nonlinear function was exactly matched to that of a CNN unit, we would have troubles finding it if we were to forward the input image at the wrong input resolution (in terms of pixels per visual angle). It is oftentimes difficult to predict *a priori* the optimal resolution at which a certain layer extracts the right nonlinearities that best match our responses, especially when the receptive field sizes of neurons are difficult to estimate for higher visual areas, and when recording beyond the foveal region of the visual field. We thus treated the input resolution as a hyperparameter that we cross-validate on the validation set. This facilitates removing the confound between the degree of nonlinearity and receptive field growth when trying to establish hierarchical correspondence between CNN layers and the biological visual system.

**Figure S2.**
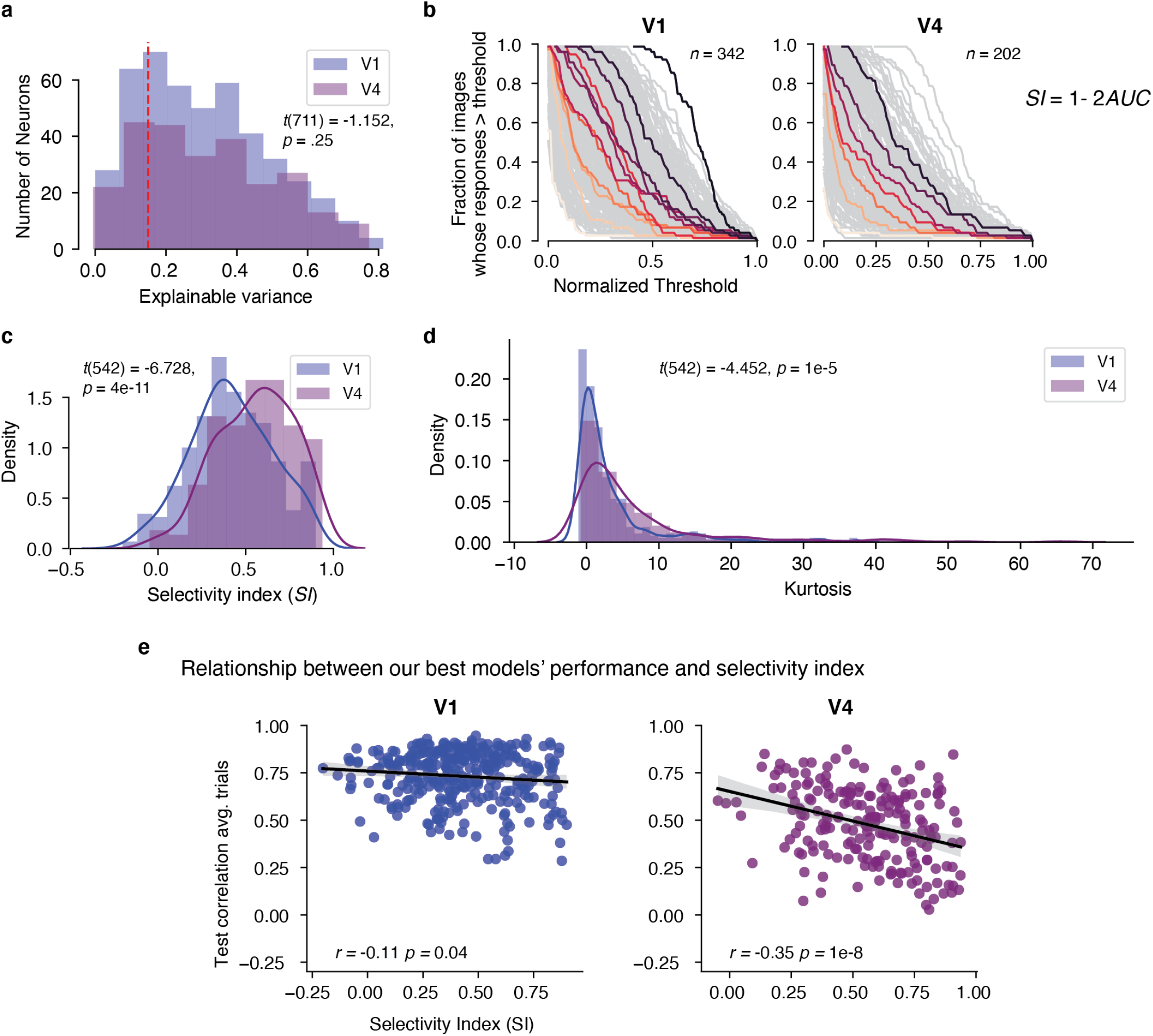
Response properties. **a**, Explainable variance (Eq. 3) distribution of V1 (*n* = 458) and V4 (*n* = 255) neurons. Red line shows the threshold (0.15) we chose to filter unreliable neurons from our test evaluations. **b**, Curves of the fraction of images evoking responses larger than a threshold vs. threshold value (normalized). We selected 100 evenly spaced thresholds between minimum and maximum value of the responses. Results for 342 neurons in V1 (left) and 202 neurons in V4 (right) show each neuron’s curve (gray). A small sample of curves were colored for clarity. We then computed for each neuron the selectivity index (SI) [62] as 1 *−* 2*AUC* where the *AUC* is the area under the curve. **c**, Density distribution of selectivity indices in V1 and V4. Two-sided *t*-test shows that means are different between areas. **d**, Density distribution of kurtosis statistic computed for each neuron over the test images in V1 and V4. Two-sided *t*-test shows that means are different between areas, highlighting increase sparsity in V4. **e**, We evaluated how well SI correlates with the predictive performance of our best model on each area (using features of Robust Resnet50) and found that selectivity index only weakly explains V1 and V4 test performance.

**Figure S3.**
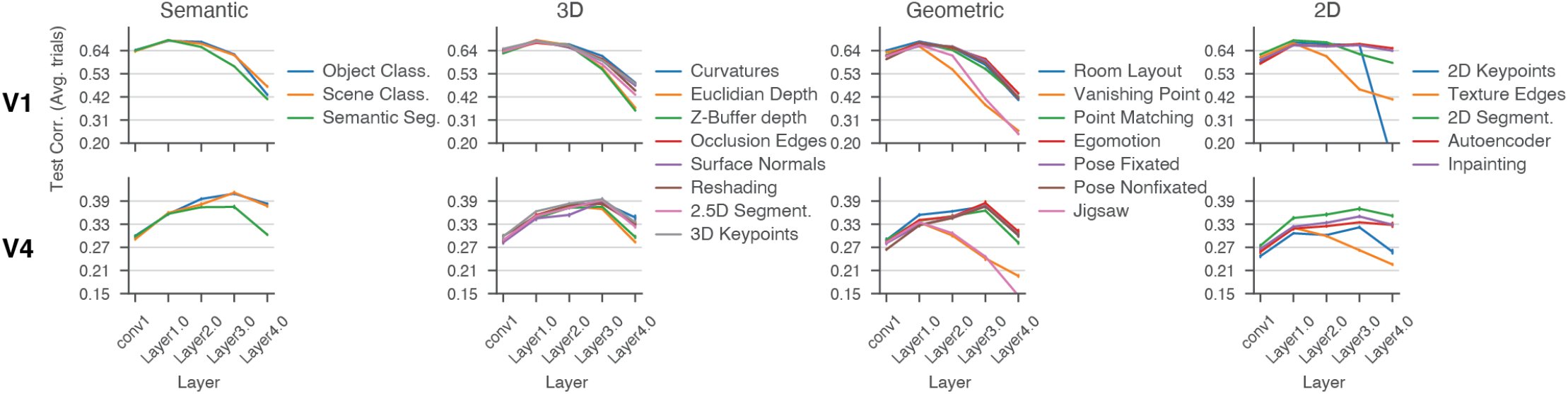
Individual task-model performances on V1 (upper row) and V4 (bottom row) as a function of network layer organized in columns by the task-clusters [15]. The task-model labels are shared between V1 and V4, and placed to the right of each column. Each line represents the average performance over seeds of the mean performance over neurons of the best task-model configuration in the validation set. That means that these lines represent the test set performance after pooling over input scales, and hyper-parameters (i.e. regularization penalty). Bars represent 95% confidence intervals of 1 s.e. of the mean for five seeds. We measured performance as the average test score over single units (*n_V_* _1_ = 458, *n_V_* _4_ = 255) calculated as the correlation between model predictions and mean responses over repetitions.

**Figure S4.**
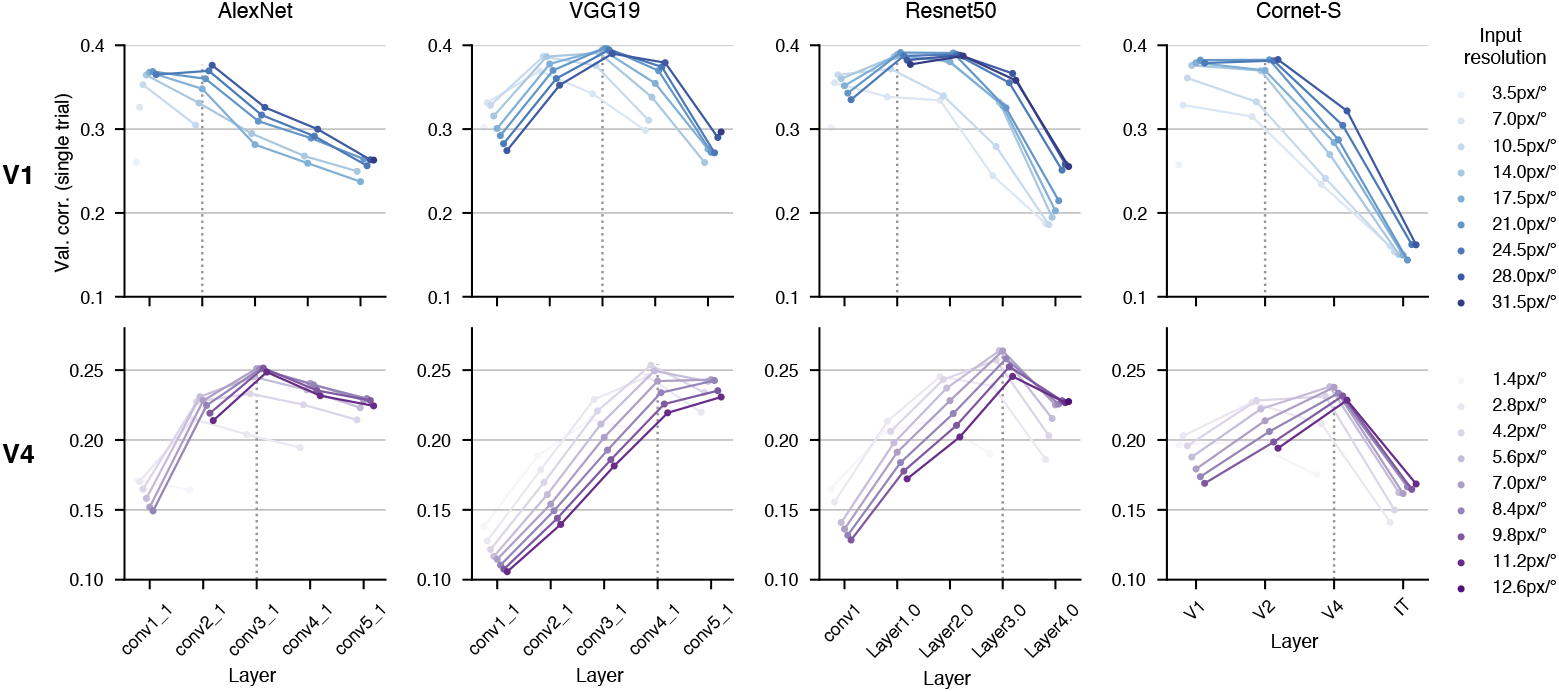
Single-trial correlation performance of ImageNet-based models at multiple input resolutions on the validation set of V1 (upper row) and V4 (bottom row). We considered four popular CNNs with different architectures, pretrained on ImageNet (columns; from left to right: AlexNet [32], VGG-19 [31], Resnet50 [18], Cornet-S [33]). For each network, we built neural predictive models that use features from multiple layers (x axis) that span the depth of the network. Each dot in the plot is the average over seeds of the best model configuration pooled over regularization parameters (see Methods). Assigning a layer to a brain area depends on the input scale – the peak of the curves shifts across input resolutions. Moreover, optimizing the layer using the wrong input resolution may lead to sub-optimal performance (Supplementary Fig. S1). We found that all of these models reveal a hierarchical ordering of nonlinear computations in the two areas, even when we account for input scale – V1 is predicted always by an earlier layer than V4 (dotted vertical lines represent the most predictive layer over scales).

**Figure S5.**
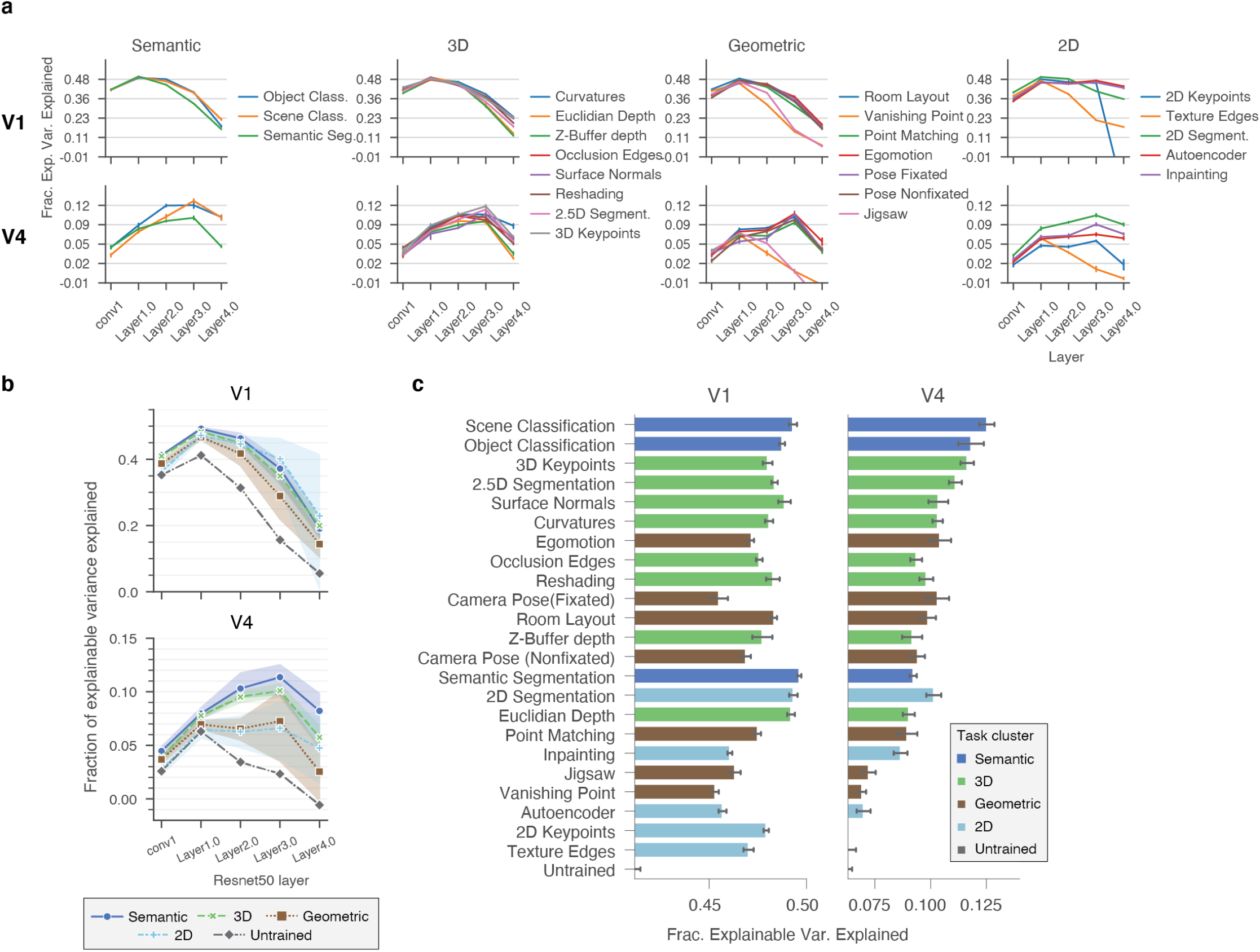
Task-model performances in terms of fraction of explainable variance explained (FEVE) **a**, Individual task-model performances on V1 (upper row) and V4 (bottom row) as a function of network layer organized in columns by the task-clusters [15]. Equivalent to Fig. S3 but performance is measured in terms of FEVE. **b**, Comparison of diverse task-driven models on V1 and V4 measured in FEVE (see Fig. 2a-b). **c**, Tasks performances in terms of FEVE after optimizing over layers and hyperparameters on the validation set ordered as in Fig. 2 c.

**Figure S6.**
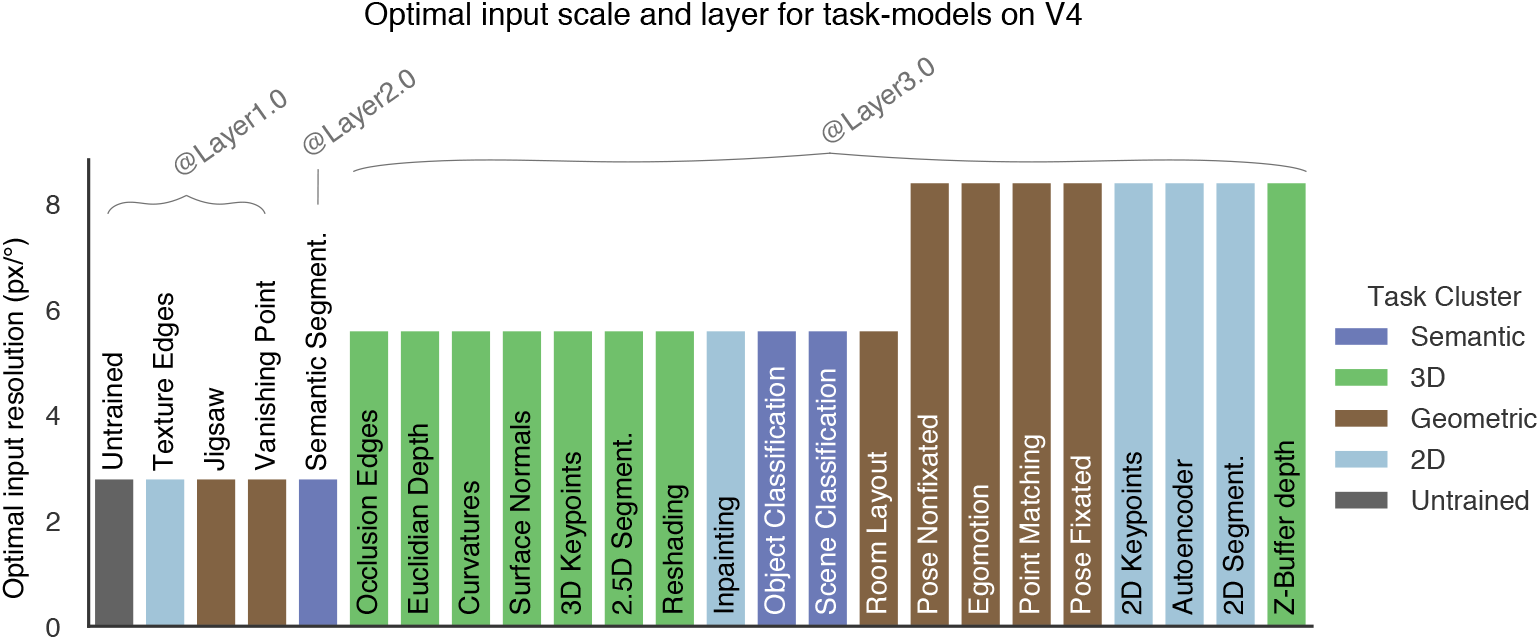
Optimal input scale and layers for task-models on V4. In contrast to area V1 where the optimal layer and scale was shared among all task-models (Layer1.0 and 21px*/^◦^*), there was variability of the optimal layer in the V4 task-models. In some models, including the untrained network, Layer1.0 with a low input resolution was optimal. The top performing models, including the two semantic classification, and most of 3D tasks (see Fig. 2c) chose an intermediate resolution (*∼* 5.6px*/^◦^*) at Layer3.0. Interestingly, most geometric and 2D tasks yielded optimal performances at the same layer, but at a higher resolution.

**Figure S7.**
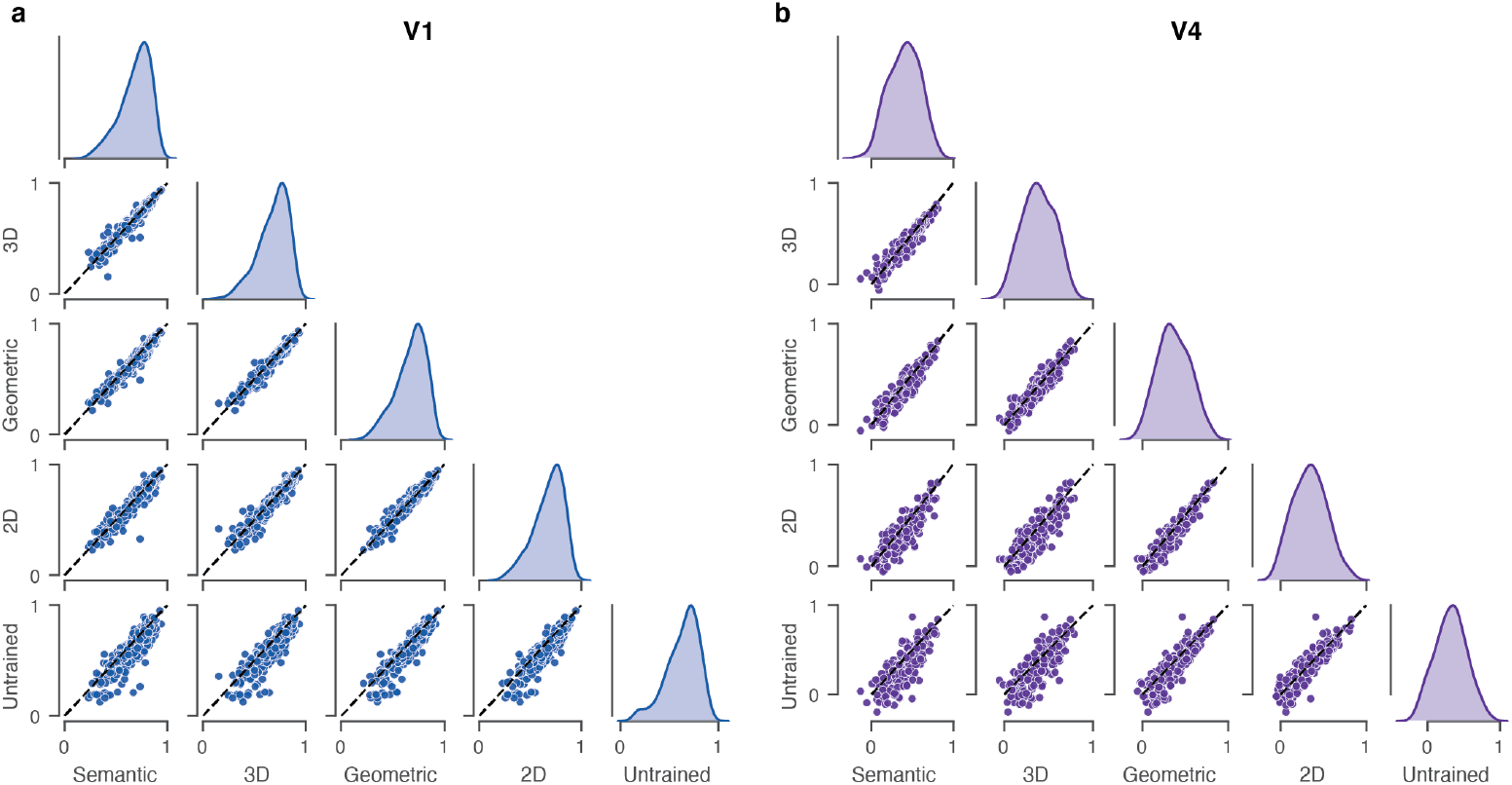
Comparison of the average task-cluster performance on single-neurons in V1. (**a**) and V4 (**b**). The dotted line in each pairwise comparison represents the identity and the panels in the main diagonal shows the performance distribution of each task-cluster. A pairwise Wilcoxon signed rank test reveal that the differences between task-clusters were significant (see Fig. 2d,e).

## References

1. Pasupathy A, Popovkina DV, Kim T. Visual functions of primate area V4. Annual Review of Vision Science. 2020;6:363–385.

2. Pasupathy A, Connor CE. Shape representation in area V4: position-specific tuning for boundary conformation. Journal of neurophysiology. 2001;86(5):2505–2519.

3. Bushnell BN, Harding PJ, Kosai Y, Bair W, Pasupathy A. Equiluminance cells in visual cortical area V4. Journal of Neuroscience. 2011;31(35):12398–12412.

4. Kim T, Bair W, Pasupathy A. Neural coding for shape and texture in macaque area V4. Journal of Neuroscience. 2019;39(24):4760–4774.

5. Conway BR, Moeller S, Tsao DY. Specialized color modules in macaque extrastriate cortex. Neuron. 2007;56(3):560–573.

6. Oleskiw TD, Nowack A, Pasupathy A. Joint coding of shape and blur in area V4. Nature communications. 2018;9(1):1–13.

7. Hanazawa A, Komatsu H. Influence of the direction of elemental luminance gradients on the responses of V4 cells to textured surfaces. Journal of Neuroscience. 2001;21(12):4490–4497.

8. Yamins DL, Hong H, Cadieu CF, Solomon EA, Seibert D, DiCarlo JJ. Performance-optimized hierarchical models predict neural responses in higher visual cortex. Proceedings of the national academy of sciences. 2014;111(23):8619–8624.

9. Pospisil DA, Pasupathy A, Bair W. ’Artiphysiology’reveals V4-like shape tuning in a deep network trained for image classification. Elife. 2018;7:e38242.

10. Bashivan P, Kar K, DiCarlo JJ. Neural population control via deep image synthesis. Science. 2019;364(6439):eaav9436.

11. Willeke KF, Restivo K, Franke K, Nix AF, Cadena SA, Shinn T, et al. Deep learning-driven characterization of single cell tuning in primate visual area V4 unveils topological organization. bioRxiv. 2023; p. 2023–05.

12. Wang A, Tarr M, Wehbe L. Neural taskonomy: Inferring the similarity of task-derived representations from brain activity. Advances in Neural Information Processing Systems. 2019;32.

13. Dwivedi K, Roig G. Representation similarity analysis for efficient task taxonomy &amp; transfer learning. In: Proceedings of the IEEE/CVF Conference on Computer Vision and Pattern Recognition; 2019. p. 12387–12396.

14. Conwell C, Prince JS, Alvarez GA, Konkle T. What can 5.17 billion regression fits tell us about artificial models of the human visual system? In: SVRHM 2021 Workshop@ NeurIPS; 2021.

15. Zamir AR, Sax A, Shen W, Guibas LJ, Malik J, Savarese S. Taskonomy: Disentangling task transfer learning. In: Proceedings of the IEEE conference on computer vision and pattern recognition; 2018. p. 3712–3722.

16. Srinath R, Emonds A, Wang Q, Lempel AA, Dunn-Weiss E, Connor CE, et al. Early emergence of solid shape coding in natural and deep network vision. Current Biology. 2021;31(1):51–65.

17. Lurz K, Bashiri M, Willeke K, Jagadish A, Wang E, Walker E, et al. Generalization in data-driven models of primary visual cortex. In: Ninth International Conference on Learning Representations (ICLR 2021); 2021.

18. He K, Zhang X, Ren S, Sun J. Deep residual learning for image recognition. In: Proceedings of the IEEE conference on computer vision and pattern recognition; 2016. p. 770–778.

19. Denfield GH, Ecker AS, Shinn TJ, Bethge M, Tolias AS. Attentional fluctuations induce shared variability in macaque primary visual cortex. Nature communications. 2018;9(1):1–14.

20. Cadena SA, Denfield GH, Walker EY, Gatys LA, Tolias AS, Bethge M, et al. Deep convolutional models improve predictions of macaque V1 responses to natural images. PLoS computational biology. 2019;15(4):e1006897.

21. Russakovsky O, Deng J, Su H, Krause J, Satheesh S, Ma S, et al. Imagenet large scale visual recognition challenge. International journal of computer vision. 2015;115(3):211–252.

22. Yamins DL, DiCarlo JJ. Using goal-driven deep learning models to understand sensory cortex. Nature neuroscience. 2016;19(3):356–365.

23. Felleman DJ, Van Essen DC. Distributed hierarchical processing in the primate cerebral cortex. Cerebral cortex (New York, NY: 1991). 1991;1(1):1–47.

24. Cadena SA, Sinz FH, Muhammad T, Froudarakis E, Cobos E, Walker EY, et al. How well do deep neural networks trained on object recognition characterize the mouse visual system? In: Advances in Neural Information Processing (NeurIPS) Neuro-AI Workshop; 2019.Available from: https://openreview.net/forum?id=rkxcXmtUUS.

25. Storrs KR, Kietzmann TC, Walther A, Mehrer J, Kriegeskorte N. Diverse deep neural networks all predict human inferior temporal cortex well, after training and fitting. Journal of Cognitive Neuroscience. 2021;33(10):2044–2064.

26. Dapello J, Marques T, Schrimpf M, Geiger F, Cox D, DiCarlo JJ. Simulating a primary visual cortex at the front of CNNs improves robustness to image perturbations. Advances in Neural Information Processing Systems. 2020;33:13073–13087.

27. Steder B, Rusu RB, Konolige K, Burgard W. NARF: 3D range image features for object recognition. In: Workshop on Defining and Solving Realistic Perception Problems in Personal Robotics at the IEEE/RSJ Int. Conf. on Intelligent Robots and Systems (IROS). vol. 44; 2010. p. 2.

28. Chen T, Kornblith S, Norouzi M, Hinton G. A simple framework for contrastive learning of visual representations. In: International conference on machine learning. PMLR; 2020. p. 1597–1607.

29. Geirhos R, Rubisch P, Michaelis C, Bethge M, Wichmann FA, Brendel W. ImageNet-trained CNNs are biased towards texture; increasing shape bias improves accuracy and robustness. In: International Conference on Learning Representations; 2019.Available from: https://openreview.net/forum?id=Bygh9j09KX.

30. Salman H, Ilyas A, Engstrom L, Kapoor A, Madry A. Do adversarially robust imagenet models transfer better? Advances in Neural Information Processing Systems. 2020;33:3533–3545.

31. Simonyan K, Zisserman A. Very deep convolutional networks for large-scale image recognition. arXiv preprint arXiv:14091556. 2014;.

32. Krizhevsky A, Sutskever I, Hinton GE. Imagenet classification with deep convolutional neural networks. Advances in neural information processing systems. 2012;25:1097–1105.

33. Kubilius J, Schrimpf M, Nayebi A, Bear D, Yamins DL, DiCarlo JJ. Cornet: Modeling the neural mechanisms of core object recognition. BioRxiv. 2018; p. 408385.

34. Zhuang C, Yan S, Nayebi A, Schrimpf M, Frank MC, DiCarlo JJ, et al. Unsupervised neural network models of the ventral visual stream. Proceedings of the National Academy of Sciences. 2021;118(3).

35. Kong NC, Margalit E, Gardner JL, Norcia AM. Increasing neural network robustness improves match to macaque V1 eigenspectrum, spatial frequency preference and predictivity. PLOS Computational Biology. 2022;18(1):e1009739.

36. Schrimpf M, Kubilius J, Hong H, Majaj NJ, Rajalingham R, Issa EB, et al. Brain-Score: Which Artificial Neural Network for Object Recognition is most Brain-Like? bioRxiv preprint. 2018;.

37. Güçlü U, van Gerven MA. Deep neural networks reveal a gradient in the complexity of neural representations across the ventral stream. Journal of Neuroscience. 2015;35(27):10005–10014.

38. Hubel DH, Wiesel TN. Receptive fields, binocular interaction and functional architecture in the cat’s visual cortex. The Journal of physiology. 1962;160(1):106–154.

39. Bay H, Ess A, Tuytelaars T, Van Gool L. Speeded-up robust features (SURF). Computer vision and image understanding. 2008;110(3):346–359.

40. Willmore B, Prenger RJ, Wu MCK, Gallant JL. The berkeley wavelet transform: a biologically inspired orthogonal wavelet transform. Neural computation. 2008;20(6):1537–1564.

41. Carandini M, Heeger DJ. Normalization as a canonical neural computation. Nature Reviews Neuroscience. 2012;13(1):51–62.

42. Burg MF, Cadena SA, Denfield GH, Walker EY, Tolias AS, Bethge M, et al. Learning divisive normalization in primary visual cortex. PLOS Computational Biology. 2021;17(6):e1009028.

43. Ronneberger O, Fischer P, Brox T. U-net: Convolutional networks for biomedical image segmentation. In: Medical Image Computing and Computer-Assisted Intervention–MICCAI 2015: 18th International Conference, Munich, Germany, October 5-9, 2015, Proceedings, Part III 18. Springer; 2015. p. 234–241.

44. Lin TY, Dollár P, Girshick R, He K, Hariharan B, Belongie S. Feature pyramid networks for object detection. In: Proceedings of the IEEE conference on computer vision and pattern recognition; 2017. p. 2117–2125.

45. He K, Gkioxari G, Dollár P, Girshick R. Mask r-cnn. In: Proceedings of the IEEE international conference on computer vision; 2017. p. 2961–2969.

46. El-Shamayleh Y, Pasupathy A. Contour curvature as an invariant code for objects in visual area V4. Journal of Neuroscience. 2016;36(20):5532–5543.

47. Sanghavi S, Jozwik KM, DiCarlo JJ. SanghaviJozwik2020; 2021. Available from: osf.io/fhy36.

48. Golan T, Raju PC, Kriegeskorte N. Controversial stimuli: Pitting neural networks against each other as models of human cognition. Proceedings of the National Academy of Sciences. 2020;117(47):29330–29337.

49. Dwivedi K, Bonner MF, Cichy RM, Roig G. Unveiling functions of the visual cortex using task-specific deep neural networks. PLoS computational biology. 2021;17(8):e1009267.

50. Mehrer J, Spoerer CJ, Kriegeskorte N, Kietzmann TC. Individual differences among deep neural network models. Nature communications. 2020;11(1):5725.

51. Geirhos R, Narayanappa K, Mitzkus B, Bethge M, Wichmann FA, Brendel W. On the surprising similarities between supervised and self-supervised models. arXiv preprint arXiv:201008377. 2020;.

52. Safarani S, Nix A, Willeke K, Cadena S, Restivo K, Denfield G, et al. Towards robust vision by multi-task learning on monkey visual cortex. Advances in Neural Information Processing Systems. 2021;34:739–751.

53. Geirhos R, Narayanappa K, Mitzkus B, Thieringer T, Bethge M, Wichmann FA, et al. The bittersweet lesson: data-rich models narrow the behavioural gap to human vision. Journal of Vision. 2022;22(14):3273–3273.

54. Kietzmann TC, Spoerer CJ, Sörensen LK, Cichy RM, Hauk O, Kriegeskorte N. Recurrence is required to capture the representational dynamics of the human visual system. Proceedings of the National Academy of Sciences. 2019;116(43):21854–21863.

55. Walker EY, Sinz FH, Cobos E, Muhammad T, Froudarakis E, Fahey PG, et al. Inception loops discover what excites neurons most using deep predictive models. Nature neuroscience. 2019;22(12):2060–2065.

56. Ding Z, Tran DT, Ponder K, Cobos E, Ding Z, Fahey PG, et al. Bipartite invariance in mouse primary visual cortex. bioRxiv. 2023;.

57. Cadena SA, Weis MA, Gatys LA, Bethge M, Ecker AS. Diverse feature visualizations reveal invariances in early layers of deep neural networks. In: Proceedings of the European Conference on Computer Vision (ECCV); 2018. p. 217–232.

58. Calabrese A, Paninski L. Kalman filter mixture model for spike sorting of non-stationary data. Journal of neuroscience methods. 2011;196(1):159–169.

59. Shan KQ, Lubenov EV, Siapas AG. Model-based spike sorting with a mixture of drifting t-distributions. Journal of neuroscience methods. 2017;288:82–98.

60. Ecker AS, Berens P, Cotton RJ, Subramaniyan M, Denfield GH, Cadwell CR, et al. State dependence of noise correlations in macaque primary visual cortex. Neuron. 2014;82(1):235–248.

61. Deng J, Dong W, Socher R, Li LJ, Li K, Fei-Fei L. Imagenet: A large-scale hierarchical image database. In: 2009 IEEE conference on computer vision and pattern recognition. Ieee; 2009. p. 248–255.

62. Quiroga RQ, Reddy L, Koch C, Fried I. Decoding visual inputs from multiple neurons in the human temporal lobe. Journal of neurophysiology. 2007;98(4):1997–2007.

63. Ioffe S, Szegedy C. Batch normalization: Accelerating deep network training by reducing internal covariate shift. In: International conference on machine learning. PMLR; 2015. p. 448–456.

64. Klindt DA, Ecker AS, Euler T, Bethge M. Neural system identification for large populations separating” what” and” where”. arXiv preprint arXiv:171102653. 2017;.

65. Sinz FH, Ecker AS, Fahey PG, Walker EY, Cobos E, Froudarakis E, et al. Stimulus domain transfer in recurrent models for large scale cortical population prediction on video. BioRxiv. 2018; p. 452672.

66. Clevert DA, Unterthiner T, Hochreiter S. Fast and accurate deep network learning by exponential linear units (elus). arXiv preprint arXiv:151107289. 2015;.

67. Kingma DP, Ba J. Adam: A method for stochastic optimization. arXiv preprint arXiv:14126980. 2014;.

68. Shi J, Malik J. Normalized cuts and image segmentation. IEEE Transactions on pattern analysis and machine intelligence. 2000;22(8):888–905.

69. Zhou B, Lapedriza A, Xiao J, Torralba A, Oliva A. Learning deep features for scene recognition using places database. Advances in neural information processing systems. 2014;27.

70. Lin TY, Maire M, Belongie S, Hays J, Perona P, Ramanan D, et al. Microsoft coco: Common objects in context. In: European conference on computer vision. Springer; 2014. p. 740–755.

71. Paszke A, Gross S, Massa F, Lerer A, Bradbury J, Chanan G, et al. PyTorch: An Imperative Style, High-Performance Deep Learning Library. In: Wallach H, Larochelle H, Beygelzimer A, d’Alché-Buc F, Fox E, Garnett R, editors. Advances in Neural Information Processing Systems 32. Curran Associates, Inc.; 2019. p. 8024–8035. Available from: http://papers.neurips.cc/paper/9015-pytorch-an-imperative-style-high-performance-deep-learning-library.pdf.

72. Harris CR, Millman KJ, van der Walt SJ, Gommers R, Virtanen P, Cournapeau D, et al. Array programming with NumPy. Nature. 2020;585(7825):357–362. doi:10.1038/s41586-020-2649-2.

73. Van der Walt S, Schönberger JL, Nunez-Iglesias J, Boulogne F, Warner JD, Yager N, et al. scikit-image: image processing in Python. PeerJ. 2014;2:e453.

74. Hunter JD. Matplotlib: A 2D graphics environment. Computing in Science & Engineering. 2007;9(3):90–95. doi:10.1109/MCSE.2007.55.

75. Waskom M, Botvinnik O, Ostblom J, Gelbart M, Lukauskas S, Hobson P, et al. mwaskom/seaborn: v0. 10.1 (April 2020). zenodo. 2020;.

76. Yatsenko D, Reimer J, Ecker AS, Walker EY, Sinz F, Berens P, et al. DataJoint: managing big scientific data using MATLAB or Python. BioRxiv. 2015; p. 031658.

77. Kluyver T, Ragan-Kelley B, Pérez F, Granger B, Bussonnier M, Frederic J, et al. Jupyter Notebooks – a publishing format for reproducible computational workflows. In: Loizides F, Schmidt B, editors. Positioning and Power in Academic Publishing: Players, Agents and Agendas. IOS Press; 2016. p. 87 – 90.

78. Merkel D. Docker: lightweight linux containers for consistent development and deployment. Linux journal. 2014;2014(239):2.

